# A network-based analysis of signal use during approach interactions across sexes in chacma baboons (*Papio ursinus griseipes*)

**DOI:** 10.1101/2023.02.04.527103

**Authors:** Jana Muschinski, Alexander Mielke, Susana Carvalho

## Abstract

Greetings in primates fulfil important functions including navigation of rank, maintenance of social relationships, and potentially establishing coalition partnerships. *Papio* makes a particularly valuable study genus as baboons show variation in greeting, male-male cooperation, philopatry, and social system. However, baboon greeting research has largely focused on male-male interactions, with female approach behaviour neglected except in relation to friendships and grunting. Most if not all signals seen in male-male greetings are also present in approaches between other sex combinations. To understand these signals further, their use in all sex combinations should be explored. We investigated approaches between male and female adult chacma baboons (*Papio ursinus griseipes*), the only savannah baboon reportedly lacking male-male cooperation, recorded in Gorongosa National Park, Mozambique. We compared male-male greetings with those of other baboon species, identified network clusters of co-occurring signals, and compared male and female approaches more broadly. Male-male approaches were similar to those in other baboon species. We identified several predictable signal combinations, ear-flattening with lip-smacking being a particularly strong signal of benign intent across sexes. Further research comparing greeting across sex combinations and species will help disentangle links between risk, cooperation, and greeting behaviour.

## 1 Introduction

Baboon greeting behaviour, studied extensively for over five decades, has recently received renewed interest due to the study of Guinea baboons, which exhibit highly physical and frequent male-male greetings compared to other baboon species [1]. Greetings between adult male baboons, which are potentially high-risk due to close physical contact, may be related to the formation and testing of coalitions, and may have deeper implications for the evolution of ritualized behaviour and cooperation, male tolerance, and sexual dimorphism across *Papio* [2]. In baboons, “greeting” refers to male-male approaches which involve some combination of a swaggering gait, ear-flattening, lip-smacking, presenting, and, depending upon species, physical contact behaviours including mounting, hip grasping, and genital touching [1–4]. Any further use of the term “greeting” will refer only to these types of male-male encounters, in line with preceding literature. We will use the terms “approaches” and “approach behaviour” more broadly because we address all sex combinations and wish to avoid conflation between context and potential function.

Approach behaviour in other sex combinations remains relatively unexplored, with the exception of female grunting behaviour and its relevance for “friendships” [5–8]. Many behaviours described in male greetings are also present in approaches involving females, including presenting, hindquarter touches, mounting, ear-flattening, and lip-smacking, yet sex combinations are usually analysed separately and rarely compared [9,10]. Studies which have included females have focused on single behaviours such as hindquarter presentations [9] or vocalizations [11]. Furthermore, existing greeting literature has focused on the presence or absence of individual behaviours or signals, rather than the usage and meaning of signal combinations. Together these issues have resulted in a poor understanding of how specific these signals and their combinations are to male-male encounters and to certain *Papio* species. Failing to understand these nuances makes it difficult if not impossible to address larger evolutionary and communication questions.

“Greeting” in itself is a problematic term as it implies the function of “saying hello”, when it is often difficult to differentiate between signals performed upon arrival that may have specific meanings (e.g., initiate grooming) and those specific to the function of greeting [12]. Across non-human primates, signals used during encounters between individuals or during merging of groups have been studied in chimpanzees, mantled howler monkeys, baboons, Tonkean macaques, sooty mangabeys, grey mouse lemurs, and black-and-white colobus monkeys, among others [11–24]. Much of the literature focuses on interactions between approaching or passing individuals, with a variety of potential underlying goals including affiliation, infant access, prevention of aggression, or reconciliation [17,20,23–25]. Accordingly, the effects of rank, social relationships, familial ties, and recent interaction history differ across studies. A second subset focuses on interactions after separation, whether these be group-level, for example the reunification of two subgroups, or individual interactions following fusion events [14,21,23]. The collections of signals used in an encounter can be visual, vocal, tactile, or bi/multi-modal [12].

In baboons, higher rates of physical greeting, particularly high-risk physical greetings, are correlated with increased male coalitionary behaviour and spatial tolerance, with chacma baboons being the non-coalition forming outlier [2]. It has been suggested that baboon greeting may be an example of ritualized behaviour and that there is a link between presence and intensity of male-male greetings and social system, degree of sexual dimorphism, and male-male competition [2,26–28]. In humans, rituals may enhance social cohesion, reduce competition, and enforce adherence to normative values [2,29–31]. The fossil record indicates that strong evolutionary parallels exist between papionines and hominins, having faced similar adaptive challenges during their parallel periods of expansion and dispersal across Africa during the Pliocene [32]. It is possible that similar adaptations, including those relating to cooperation and social cohesion, could have allowed both clades to succeed in novel and changing environments.

The high level of variation in social structure, male-male relationships, cooperation, sexual dimorphism, and socioecology in the *Papio* genus provides an ideal natural experiment to study relationships between these factors. Six species of baboon currently range through a variety of environments across Africa and the Arabian Peninsula (*P. hamadryas, P. papio, P. anubis, P. cynocephalus, P. ursinus, P. kindae*), with several hybridization zones [33,34]. Their environments vary drastically between and within species, which may influence signalling repertoire and frequency [35,36]. While sexual dimorphism across *Papio* is relatively high in comparison to other genera, chacma baboons are noticeably more dimorphic in canine height than the other *Papio* species and hover at the high end of spectrum of male to female body mass ratios in the genus (see table 1.1) [37–39]. Understanding how males navigate interpersonal relationships, particularly in species with intense competition, is critical for studying relationships between behaviour, male competition, social structure, and the evolution of sexual dimorphism. Differing levels of sexual dimorphism may alter perceived risk levels in approaches between different sex combinations across the *Papio* species, resulting in differences in signal use. Conversely, an improved understanding of communication in close-range interactions across sex combinations may provide further insight into how such interactions may shape and facilitate relationships which in turn affect reproductive success.

**Table 1.1:** Greeting behaviour, philopatry type, and sexual dimorphism across *Papio*

| Species | Greeting exhibited | Physical contact | Philopatry Type | M-M Coalitions | Canine Height (M:F) [37, 38, 39, 43, 80] | Body Size (M:F) [37, 38, 39, 43, 80] |
| --- | --- | --- | --- | --- | --- | --- |
| chacma baboon | Contested [10, 42, 50] | Unknown | Female [40, 41, 42, 83] | No [40, 41, 42, 83] | 3.84 | 2.01 |
| yellow baboon | Yes [9, 84] | Unknown | Female [40, 41, 83] | Yes [40, 41, 83] | 2.80 | 1.76 |
| kinda baboon | Unknown | Unknown | Female [81] | No [81] | 3.00 [81] | 1.77 [81] |
| olive baboon | Yes [46] | Rare [46] | Female [40, 41, 83] | Yes [40, 41, 83] | 2.22 | 1.53; 1.85; 1.81; 1.89 |
| hamadryas baboon | Yes [3, 47, 84] | Rare | Male [3, 47, 84] | Yes [40, 41, 83] | 2.74 | 1.84; 1.78; 1.71 |
| Guinea baboon | Yes [1, 27] | 93.40% | Male [43] | Yes [40, 41, 83] | 3.14 | 1.7 |

The four “COKY” baboons (chacma, olive, Kinda, and yellow baboons) all exhibit multi-male, multi-female groups with polygynandrous mating systems, female philopatry, and male dispersal [40]. Male coalitions have been reported consistently across all COKY baboons except the chacma baboon [40,41]. Limited male coalitionary behaviour was observed in one male-male pair of chacma baboons by Saayman [42] and reported in chacma baboons in Gorongosa National Park during hunting activity (personal communication, Susana Carvalho), suggesting under-reported variation. Unlike the COKY baboons, Guinea and hamadryas baboons exhibit multi-level hierarchical social structures. In both species, males are philopatric, remaining in their natal clan/party, while females disperse from their natal groups [43,44]. Unlike hamadryas baboons, male Guinea baboons demonstrate strong bonds with other males, with high levels of male – male tolerance and affiliative behaviours such as grooming, even between less closely related males [45].

Greetings occur in hamadryas and Guinea baboons, and at a lesser rate, yellow and olive baboons [1,9,46,47]. The most ritualized and physical of greetings are exhibited by Guinea baboons; more varieties of physical contact are exhibited and physical contact is generally more frequent, intense, and risky than in the other species [1,2,27]. Greetings take on a variety of functions in male baboons, at times dependent upon the species, and may assist with in-group identification, bond testing, and relationship reinforcement (Guinea baboons) [1,27], test potential for coalitions (olive baboons) [46], or ease tension and avoid confrontation through signalling of competitive power (hamadryas baboons) [3,47,48]. Systematic study of greeting in chacma baboons is limited (Tables 1.1 and 3.2), despite the importance of chacma baboons when studying relationships between greeting behaviour, coalitionary behaviour, and sexual dimorphism [10,41]. They are generally reported as exhibiting limited greeting behaviour, with little to no physical or high-risk (i.e., genital) contact [2,10,49]. While chacma baboons do exhibit some of the less intense greeting behaviours reported in other species, close proximity approaches by male chacma baboons, whether towards a recipient adult male or adult female, are more likely to happen without greeting behaviour than with and physical contact is rare [10,42,50]. Given their status as a potential outlier among *Papio* species, further systematised research on chacma greeting behaviour would be a valuable addition to the literature.

While most female baboon approach behaviour has been understudied, grunting is the exception [6,7,51]. Suggested functions of grunting include signalling benign intent, reinforcing social bonds, indicating high arousal, and reconciling following agonistic interactions [52,53]. Across baboon species, grunting is more common when infants are present and may be dependent on social bond strength and familial relationship [53]. Grunts may be used in a reconciliatory context and interactions following grunting are less likely to be agonistic and more likely to involve infant contact [5,51,52]. Differences in methodology between the female grunting and male greeting literatures make direct comparison between different sex combinations difficult. A more encompassing view of baboon approach behaviour would contribute significantly to our understanding of the signals used during approaches, how signals are combined, and how their use relates to the sex and goals of approacher and recipient.

### 1.1 Reconsidering baboon “greeting”

We find that there are three primary issues at play in the existing *Papio* approach literature. First, the study of greetings does not consider females and how the same signals are used in approaches between other sex combinations. Second, there is insufficient data on chacma baboon approach behaviour and limited understanding of within-species variation. Third, much of the existing approach behaviour and greeting literature in baboons focuses on the presence and absence of individual signals, rather than considering how signals are combined. This is an issue across the primate literature more broadly and makes it difficult to identify how multi-modal signalling is composed and how small differences in composition modify meaning [12,54]. In chimpanzees, for example, the likelihood of a reciprocal greeting is strongly influenced by the modality of the initial greeting [19].

Here, we study approach behaviour in male and female chacma baboons (*P. ursinus griseipes*) using video footage from Gorongosa National Park, Mozambique. Rather than focusing solely on interactions where “greeting behaviours” were exhibited, we record proximity events (any instance in which an individual comes within two meters of a conspecific after having previously been more than five meters away [similar approach as in 10]). Use of this broader criterion and video footage rather than *in situ* observation allows for investigation of why such behaviours are exhibited in some approaches, but not others, and prevents the accidental exclusion of subtle behaviours which may be missed upon first – or live – viewing. The study has three primary objectives – to provide first, a further account of male-male approach behaviour in chacma baboons and situate this within the existing literature; second, to look at how signals are used in combination using a network approach; and third, to conduct a direct comparison of the signals used in the different sex combinations, assessing how specific to male-male approaches the use of the aforementioned “greeting” signals truly are.

We primarily applied a network analysis approach to study co-occurrence of signals across varying conditions using the NetFACS package [55], originally designed for the study of complex facial signals. Taking a network approach allows us to study the relationship between signals themselves and between signals and specific conditions, providing a greater degree of insight into the structure of approach behaviour. Each signal is treated as a network node, with network edges determined by behaviour co-occurrence [55,56]. We additionally used the package’s permutation test functionality to test predictions regarding differences in behaviour prevalence between sex combinations.

Our study aims to address the following research questions and accompanying predictions.

1. ***Research Question*** How do male-male approaches in Gorongosa chacma baboons compare to the published literature? ***Prediction*** Based on the existing literature, we would expect chacma baboons to show little to no contact behaviour, particularly intense contact, in male-male approaches when compared to other baboon species.
2. ***Research Question*** Are there specific signals which happen together more than expected and do they represent different approach “types”? ***Prediction*** We expect to identify signal clusters that may be tied to specific sex combinations or be related to specific goals (e.g., gaining infant access or receiving grooming).
3. ***Research Question*** How does the combination of approacher and recipient sex influence the signals and combinations thereof exhibited during approach? Are any of the behaviours which are frequently discussed in the male-male literature specific to male-male approaches? These behaviours include lip-smacking, ear-flattening, presenting, mounting, and hindquarter touching. ***Prediction*** We predict that sex combination influences the signals expected, with identifiable male-male versus female-female signals, but that most signals will show overlap in usage. We expect the aforementioned behaviours may be more common than expected in male-male encounters.

## 2 Methods

### 2.1 Study site and population

Gorongosa National Park covers 3770 km^2^ of the Urema drainage basin in the southern end of the East African Rift System (EARS)[57–60]. The mosaic ecosystem results in high biodiversity and makes the park a unique and valuable analogue model for the environmental conditions of the EARS during important periods of human evolution [57,61,62]. The park’s baboons are usually categorized as chacma baboons (*Papio ursinus griseipes*), but the park lies within a potential hybridization zone between northern chacma baboons and southern yellow baboons [59,63]. Our study group, the Chitengo Troop, resides in the forested area surrounding the tourist site and research centre and is well habituated due to continuous exposure to humans. As of November 2019, the troop consisted of 8 resident adult males, 3 peripheral adult males, 1 subadult male, 11 adult females, 1 subadult/large juvenile female, approximately 15 small/medium juveniles, and an indeterminate number of infants. All adult and subadult baboons were identified and named by JM in 2019 and can successfully be identified *in situ* and from sufficiently high-resolution video footage. Sixty-five hours of footage were recorded opportunistically during October and November 2018 and between July and November 2019 by colleague Lucy Baehren and JM [64]. Recording focused on groups of baboons, with target group rotated throughout the day, but was not randomized as individuals had not been identified at the time of recording. Approacher identity was controlled for *post hoc*. Video recording in Gorongosa National Park was completed under permit number PNG/DSCi/C145/2019 (J. Muschinski) and PNG/DSCi/C110/2018 (L. Baehren) and was cleared by the University of Oxford Animal Welfare and Ethical Review Board (APA/1/5/ACER/10Dec2018).

### 2.2 Video coding procedure

Footage was reviewed *post hoc* and all proximity events identified. Proximity events are here defined as any instance in which one individual, the approacher, decreases the distance between themselves and the recipient from over five meters to less than two meters. The proximity event began once the approacher entered a five-meter radius of the recipient and concluded when either 1) the approacher or recipient increased the distance between each other to over five meters or 2) 30 second had passed since the approacher came within two meters of the recipient. In most cases, the approacher could be easily identified, with one individual approaching and the other remaining stationary. If both approached each other, the individual who began approaching first was considered the approacher. Where an individual approached more than one stationary individual, the recipient was defined as the individual who the recipient interacted with or signalled towards first. If neither individual was interacted with, the first individual the approacher passed was considered the recipient. Two individuals simultaneously approaching a third occurred very rarely and in such situations the individual who came within two meters of the recipient first was considered the approacher. Juveniles were not included in these analyses because only adults and subadults could be reliably identified and identification is necessary to control for potential effects of individual relationships. Events were only included if over 50% of the entire sequence could be seen. The ethogram used to collect behavioural data was modelled primarily after Dal Pesco & Fischer [1], Colmenares [65], and Silk [66] (for full ethogram see [67]). The cleaned dataset is publicly available [68]. Behavioural data was collected using BORIS version 7.10.2 [69] and data cleaning and analysis performed using Python version 3.8.5, R version 4.2.2, and NetFACS version 0.5.0.

For analyses using NetFACS (sections 2.4 and 2.5) we included only proximity events where visibility allowed for identification of the approacher’s general facial expressions (e.g., lip-smacking) and where both individuals were adults or subadults (n = 341). Future analyses will focus on outcomes of these interactions and the behaviour of the recipient; this paper focuses specifically on the description and identification of patterns or combinations of signals exhibited by the approachers. We include only actions performed by the approacher during the approach and initial interaction. The initial interaction is defined as ending once the approacher sits or begins walking away, foraging, grooming, resting, or being groomed. NetFACS, used for permutation tests and network analysis, requires presence/absence data for each signal of interest for each event and does not account for the order, intensity, count, or length of each behaviour. It compares observed probabilities to expected probabilities created using bootstrapping [55]. Prior to analysis we combined several similar behaviours which had been split too finely during ethogram creation (e.g., combining all non-contact threats into one category, combining all types of embraces, etc.) into larger behaviour groups and we excluded any behaviours which occurred in fewer than four events (1% of events) [55,70].

### 2.3 Male-male “greeting” analysis

To enable comparison with existing literature [2], all male-male proximity events were classified as either “non-greetings” or “greetings” based on the presence of any traditional “greeting behaviour” (lip-smacking, ear-flattening, continued direct gaze, physical contact). All proximity events that could be defined as “greetings” were then assessed on three criteria - presence of any physical contact (initiated by approacher or recipient), presence of intense physical contact (initiated by approacher or recipient), and reciprocation. Intense physical contact has previously been defined as genital touching, embracing, or mounting [1,27]. Greetings were scored as “reciprocal” when the approacher and recipient both perform at least one greeting behaviour [1,3]. Percentages of greetings that were physical, intense, or reciprocal were calculated in relation to the count of male-male “greetings”, rather than in relation to all male-male proximity events, to allow for direct comparison with the literature. It should be noted here that the total number of male-male greetings reported across the 65 hours of video footage cannot be directly compared to hourly rates reported elsewhere due to differences in methodology (opportunistic videography vs. focal follows). We calculated bootstrapped 95% confidence intervals using the boot R package [71].

### 2.4 Identification of signal combinations - community detection

To determine whether types of greeting can be identified without additional information (e.g., approacher sex), we applied community detection using the NetFACS package with a modularity threshold of at least 0.3 [55,72]. NetFACS community detection uses the “fast greedy” modularity optimization algorithm to determine which groups of elements co-occur more than expected [55]. We completed this analysis twice – once with the dataset as prepared, with a quite extensive ethogram in which behaviours are included independently (35 behaviours), and once with a minimal ethogram, where behaviours are collapsed into a total of 25 categories (e.g., all types of non-maternal infant contact lumped as “infant contact,” all types of non-aggressive contact lumped into one category, etc.; see [67]). Each was run with 2000 randomizations, a minimum significance of 0.05, minimum count of 17 (5% of 341 observations), and minimum probability of 0.05. Full results of the minimized ethogram analysis are discussed below.

To determine whether these signal combinations were associated with specific sex combinations or whether the combining of all sex combinations into one analysis hid sex-specific patterns, we split the dataset into the four sex combination categories (male-male, female-female, female-male, and male-female) and completed the same analysis for each subset using the minimized ethogram. Minimum count cut-offs were adjusted for the subsets sample sizes. Graphical results for analyses of the expanded ethogram and for the four sex combination subsets are included in the supplementary information.

### 2.5 A comparison of the specificity of “greeting” behaviours to male-male approaches

We compared the use of five signals identified in most male-male greeting ethograms (lip-smacking, ear-flattening, mounting, presenting, and hindquarter touches) between male-male approaches and the other sex combinations to determine whether any of these signals or their combinations are specific to male-male approaches. For these analyses we compared observed probabilities for each behaviour or combination of interest in male-male events to the expected probability calculated from a permutation of all events (randomizations = 1000) using the full ethogram with the NetFACS package. We included a random effect of approacher ID and we controlled for presence of an infant not belonging to the approacher. An alpha value of 0.01 was used to account for multiple comparisons. We repeated the same analysis across the remaining sex combinations (female-female, male-female, female-male).

## 3 Results

A total of 428 interactions were identified from 65 hours of video footage, with 341 meeting all visibility inclusion criteria (see table 3.1). The approacher was identifiable in all but 19 of the qualifying interactions and the recipient in all but 18 (in 4 interactions neither approacher nor recipient could be identified confidently). The mean number of unique signals included per approach were 2.22 (SD = 1.47) for male-male approaches, 3.14 (SD = 2.23) for female-female approaches, 2.45 (SD = 1.72) for male-female approaches, and 1.84 (SD = 1.22) for female-male approaches when using the full ethogram.

**Table 3.1:** Counts of interaction types

| Recipient | Approacher sex |  |
| --- | --- | --- |
|  | Female | Male |
| Female | 167 | 64 |
| Male | 73 | 37 |

### 3.1 Male-male approaches - how do the chacma baboons compare?

A total of 51 male-male interactions involving only adult/subadults were identified across all 65 hours of footage. Of the 51 male-male proximity events, 43 can be considered “greetings” according to criteria applied in other studies (presence of lip-smacking, ear-flattening, physical contact, or prolonged eye contact or gaze towards) [1,2,27].

While Guinea baboons are certainly exceptional in terms of physical and intense physical contact, it does not appear that chacma baboons are an outlier when compared to the other COKY baboons (see table 3.2). Approximately 16.3% of chacma male baboon greetings involved physical contact (95% confidence interval from 5.2% to 27.1%) and 9.3% intense physical contact (95-CI: 0.5% to 18%), similarly to olive and hamadryas baboons. Reciprocity was also similar in the chacma baboon sample and previously reported hamadryas studies (estimate: 72.1%, 95-CI: 58.5% to 85.6%).

**Table 3.2:**
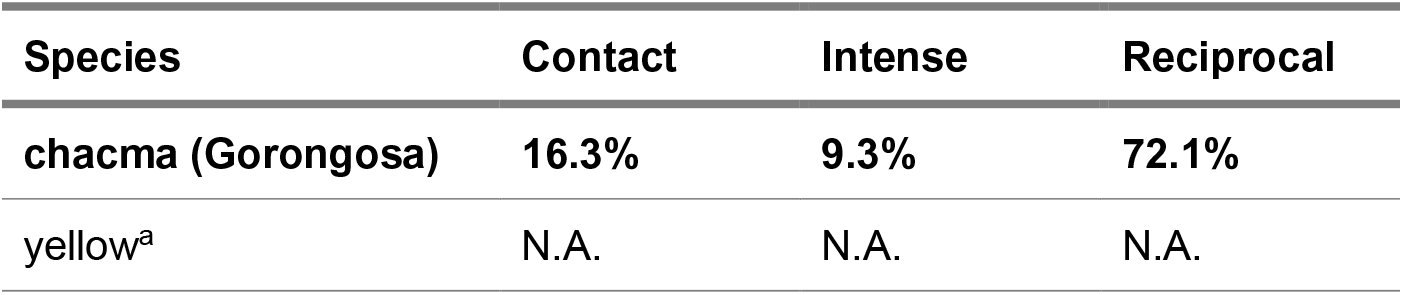
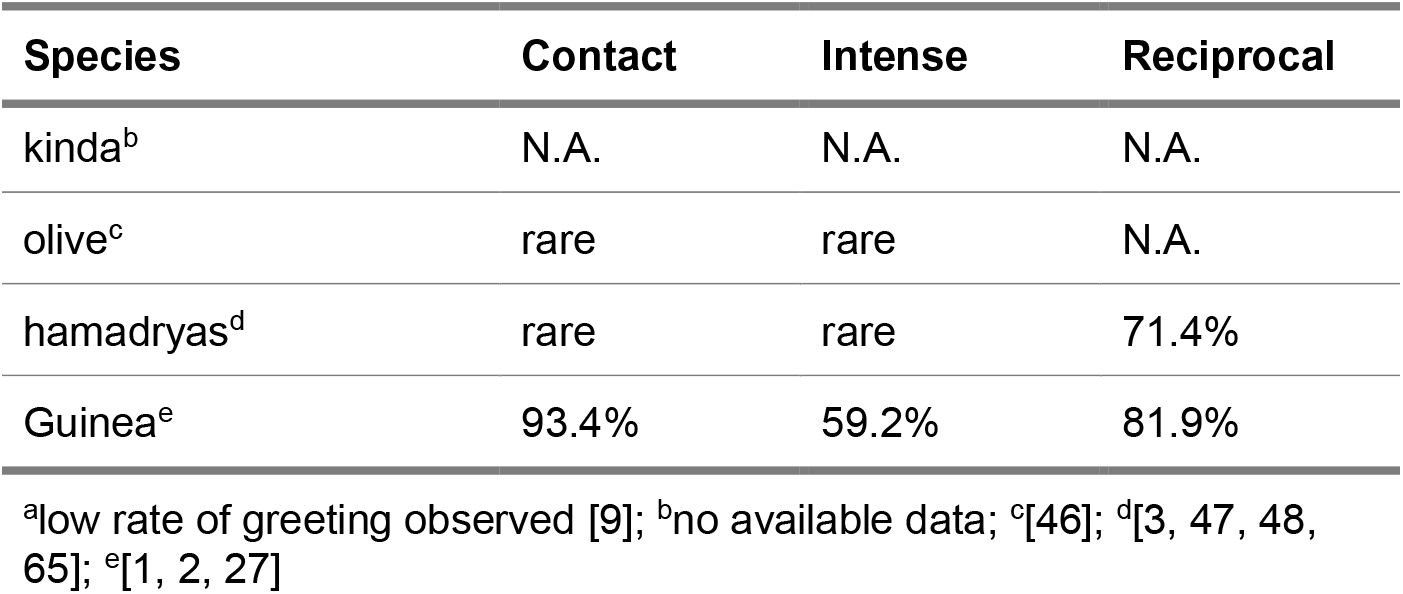
Comparison of male greetings across baboon species

| Species | Contact | Intense | Reciprocal |
| --- | --- | --- | --- |
| chacma (Gorongosa) | 16.3% | 9.3% | 72.1% |
| yellow<sup>a</sup> | N.A. | N.A. | N.A. |
| kinda<sup>b</sup> | N.A. | N.A. | N.A. |
| olive<sup>c</sup> | rare | rare | N.A. |
| hamadryas<sup>d</sup> | rare | rare | 71.4% |
| Guinea<sup>e</sup> | 93.4% | 59.2% | 81.9% |

<sup>a</sup>low rate of greeting observed [9]; <sup>b</sup>no available data; <sup>c</sup>[46]; <sup>d</sup>[3, 47, 48, 65]; <sup>e</sup>[1, 2, 27]

### 3.2 Community detection: Identification of signal combinations

Community detection was completed with the larger ethogram (35 behaviours - results in supplementary information) and again with a minimized ethogram (25 behaviours). With the minimized ethogram, community detection identified four clusters with a modularity of 0.49 (Figure 3.1). The first cluster included passing without contact and glancing toward the recipient; the second cluster contained observing the recipient, arriving (classified as an approach that ends in the individual stopping at and/or interacting with the recipient rather than diverting or passing without contact), non-aggressive physical contact, and presenting; the third consisted of observing the recipient’s infant, having physical contact with the infant, and grunting; the fourth cluster consisted of lip-smacking and ear-flattening.

**Figure 3.1:**
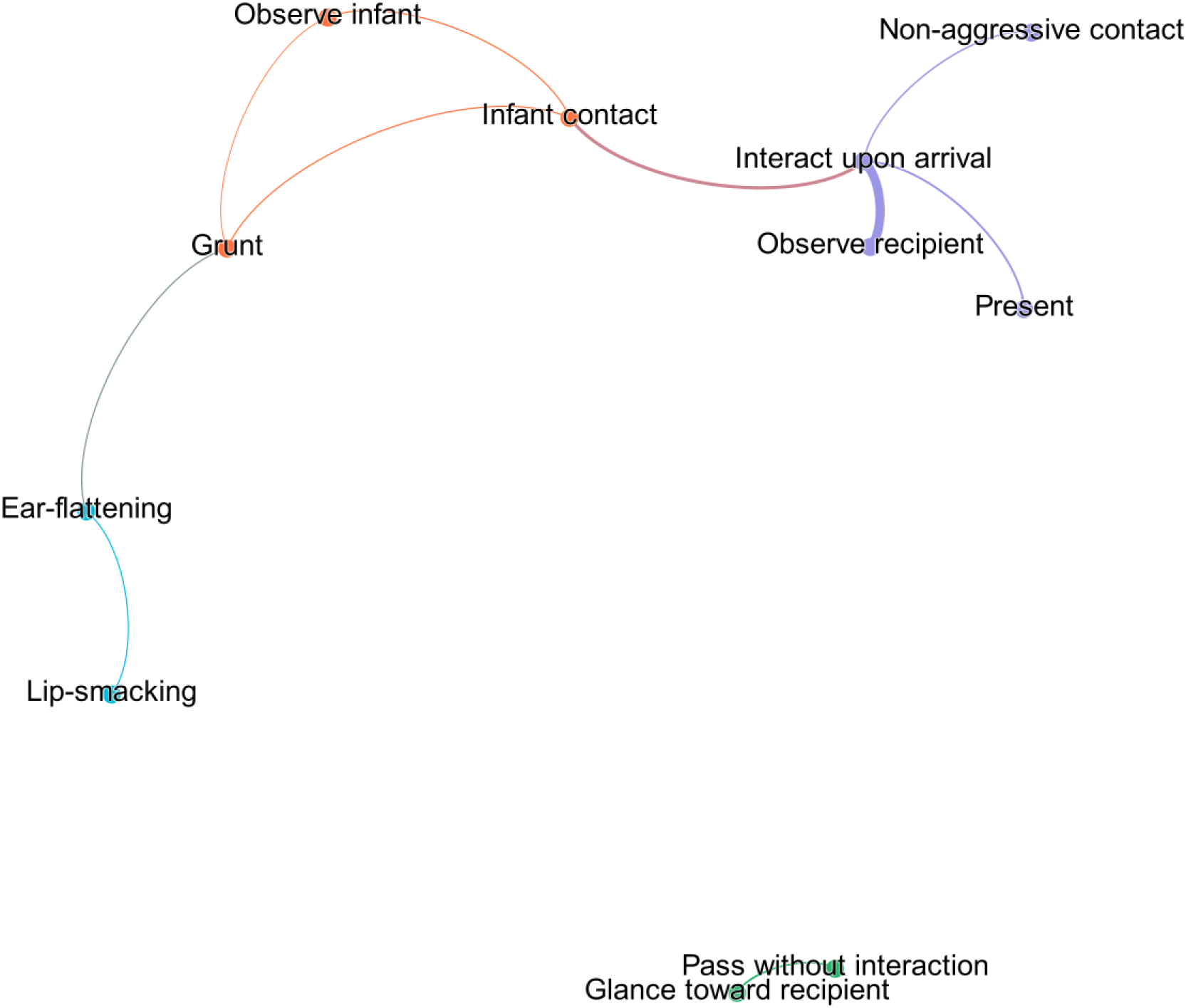
Community detection across all greetings with a minimal ethogram. Linked and coloured behaviours are detected clusters, with edges labelled with the combination’s observed probability. Figure created using Gephi.

**Figure 3.2:**
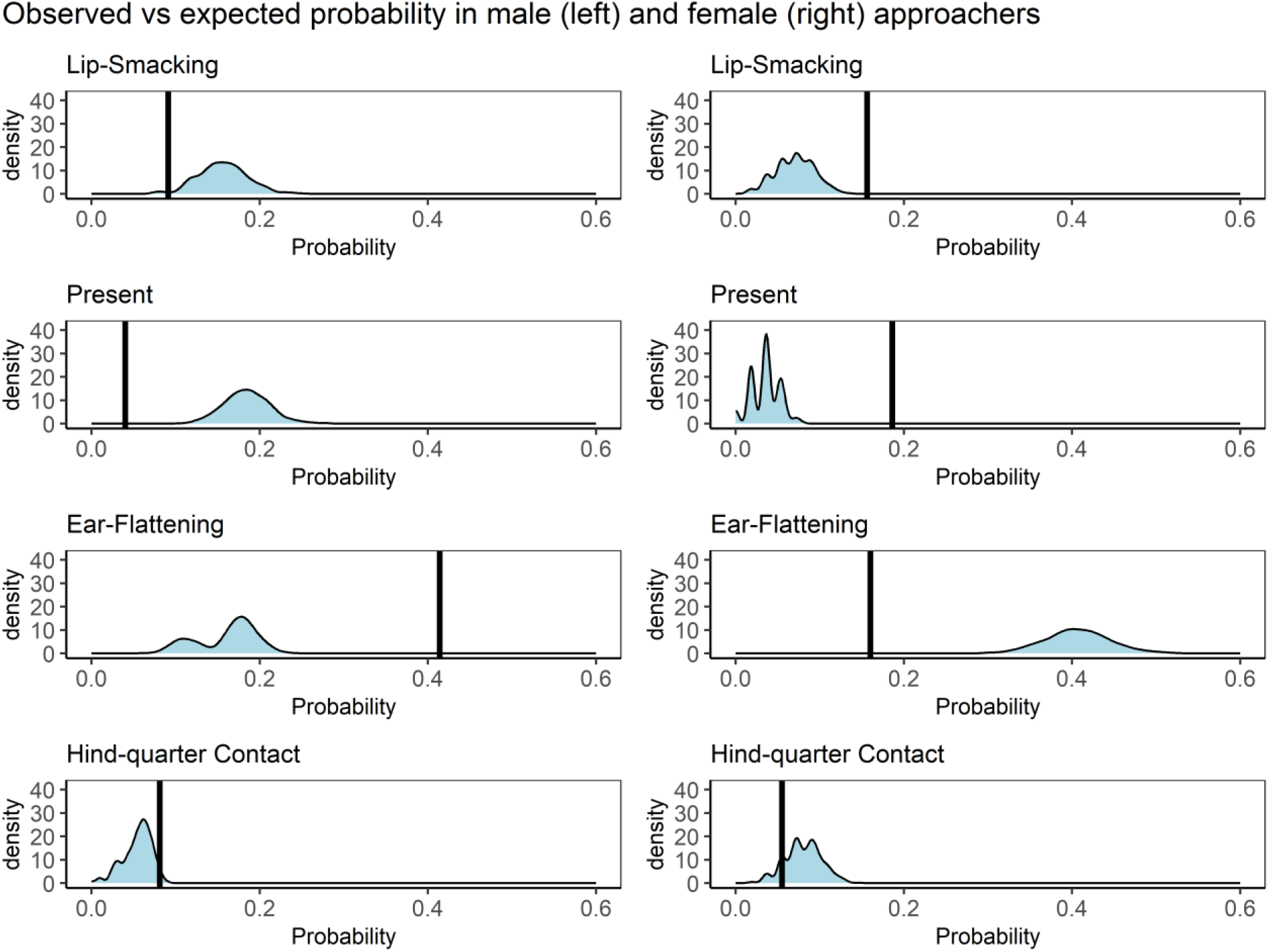
The distribution of the expected probability of four behaviours during approach based on bootstrapped samples of the opposite sex’s approach behaviours, with vertical lines representing the observed probabilities in the sex in question (male on left, female on right)

When the dataset was split into subsets by sex combination, clusters could be identified in each subset (see table 3.3, graphical results in supplementary materials). Due to the high number of female-female events, we expect community detection using only female-female events to be most similar to that using all events.

**Table 3.3:** Community detection results for sex combination subsets

| Sex Combination | Modularity | Clusters |
| --- | --- | --- |
| Female-female | 0.38 | Pass without contact and glance toward |
|  |  | Observing infant, infant contact, and interacting upon arrival |
|  |  | Lip-smacking, ear-flattening, grunting, and non-aggressive contact |
| Male-male | 0.50 | Ear-flattening, interacting upon arrival, and observing during approach |
|  |  | Grimace, present, and evasive movement |
| Female-male | 0.32 | Passing without contact and glancing toward recipient |
|  |  | Observing, presenting, pausing during approach, and interacting upon arrival |
| Male-female | 0.42 | Observing, soliciting grooming, and interacting upon arrival |
|  |  | Ear-flattening and grunting |
|  |  | Passing without contact and glancing toward recipient |

### 3.3 A comparison of signal use in male versus female approachers and across sex combinations

The five signals we aimed to compare included lip-smacking, ear-flattening, presenting, hindquarter contact, and mounting. However, due to limited occurrences and sample sizes once splitting by approacher sex and by sex combination, we did not include mounting when splitting by approacher sex or hindquarter contact and mounting when splitting by sex combination (fewer than 4 occurrences in male or male-male encounters respectively). Given the sample sizes and limited occurrences, we felt such comparisons would not be valid. However, the rarity of these behaviour indicates they may not play as significant of a role as in some other *Papio* species.

#### 3.3.1 Male versus female approachers

When comparing observed probabilities in male approaches versus a boot-strapped sample of female approaches, we found males performed ear-flattening more than expected (probability = 0.41, probability increase = 2.5, effect size = 0.25, specificity = 0.71, p < 0.001). When doing the reverse, with observed probabilities in female approaches compared to boot-strapped male approaches, we found females exhibited more lip-smacking and presenting than expected (lip-smack: probability = 0.16, probability increase = 2.2, effect size = 0.09, specificity = 0.69; present: probability = 0.19, probability increase = 5.2, effect size = 0.15, specificity = 0.88; p < 0.001 for both). Pairings of the five behaviours (for example lip-smacking with hindquarter contact) were not more common than expected for either sex. While grunting was not part of the original list of signals being tested, a comparison of all signals showed that the combination of grunting together with lip-smacking had a particularly high probability increase of 32.2 (probability = 0.08, effect size = 0.08, p < 0.001) in female versus male approaches.

#### 3.3.2 Across sex combinations

##### Male-Male

Separate analysis of ear-flattening and lip-smacking indicate both occur more frequently than expected in approaches of male-male encounters when compared to a boot-strapped sample from all sex combinations, with ear-flattening having a probability increase of a factor of 2.6 and lip-smacking one of 1.8 (ear-flattening: probability = 0.53, effect size = 0.33, specificity = 0.49; lip-smacking: probability = 0.22, effect size = 0.1, specificity = 4.47 respectively, p < 0.001 for both). While a comparison of all male approaches to all female approaches showed no significant difference in the use of the lip-smacking and ear-flattening together (p > 0.05), a comparison of specifically male-male approaches to all sex combinations did find that the observed probability of ear-flattening and lip-smacking occurring in the same approach was higher than expected in male-male encounters (probability = 0.17, probability increase = 2.3, effect size = 0.1, specificity = 0.51, p < 0.001).

##### Female-Female

In female-female encounters, lip-smacking had a higher observed probability than expected based on a boot-strapped sample of all sex combinations (probability = 0.22, probability increase = 3.4, effect size = 0.15, specificity = 0.43, p < 0.001). In combinations of size two, ear-flattening with lip-smacking had an observed probability higher than expected (probability = 0.12, probability increase = 2.4, effect size = 0.07, specificity = 0.37, p < 0.001). Though not the focus of this analysis, many combinations of infant directed behaviour, grunting, lip-smacking, and ear-flattening were also present significantly more than expected (supplementary information).

##### Female-Male

In female-male encounters, presenting had a higher observed probability than expected (probability = 0.41, probability increase = 6.3, effect size = 0.34, specificity = 0.70, p < 0.001). No size two combinations of the five relevant signals were observed more than expected.

##### Male-Female

In male-female encounters, ear-flattening was observed more than expected (probability = 0.35, probability increase = 1.68, effect size = 0.14, specificity = 0.35, p < 0.001). No size two combinations of the five relevant signals were observed more than expected.

## 4 Discussion

Our study addresses three research themes: first, how chacma baboon behaviour fits into the baboon greeting literature; second, whether signals are reliably combined; and third, in what ways approach behaviour differs across sex combinations and the implications for studying male-male events in isolation. The rates of intense contact, any contact, and reciprocal greeting in chacma baboons are similar to those observed in olive and hamadryas baboons, indicating that they may not be as extreme of an outlier as suggested by existing literature [2]. We identified four clusters during community detection, suggesting that these signal combinations occur more frequently than expected. Our analyses of male versus female approach behaviour highlight both key differences and important areas of overlap, for example in the use of the ear-flattening and lip-smacking combination.

### 4.1 Chacmas and the *Papio* genus

It has been suggested that there is little to no male-male greeting in chacma baboons and that it is far less elaborate than those of other COKY baboons [10], but this impression may stem partly from a lack of research on greeting in the species. Across only 65 hours of footage, we identified 43 “greetings,” some including hindquarter touches and even genital touching. While we cannot compare rate of greeting per hour, we found that 16% of greetings involved contact, 9% intense contact, and 72% are reciprocal. These align well with proportions seen in yellow, olive, and hamadryas baboons (see table 3.2). Dal Pesco & Fischer suggested that male-male greeting behaviour in baboons follows a geographic cline in elaboration and ritualization, with a large phylogenetic split between the southern (chacma, yellow, and kinda baboon) versus northern (olive, hamadryas, Guinea baboon) clades [2]. They point out that species where males are more spatially tolerant and affiliative also have the highest rates of greeting and most ritualized greeting behaviour, supporting suggested connections between human prosociality, larger group living, and the evolution of ritual. Guinea baboons are the noticeable outlier when it comes to male-male greetings, with a particularly high hourly rate, 93.4% of all greetings involving contact, and 59.2% involving intense physical contact; they also demonstrate high levels of male-male spatial tolerance, affiliative behaviour, and the lowest, though comparable, level of sexual dimorphism in *Papio* [1].

Existing research on greeting in chacma baboons is limited, with early studies by Saayman and Hall reporting limited presenting and contact behaviour between males [42,50]. At odds with the remaining chacma literature, Saayman does report limited male-male coalitionary behaviour, suggesting there may be within-species variation in male cooperative behaviour [42]. Kalbitzer et al. approached the study of greeting behaviour using a similar format as here, recording all approaches within one meter, and reported limited physical greeting among chacma baboons in the Moremi Game Reserve, Botswana [10]. They recorded interactions as greetings only when non-agonistic contact and non-affiliative contact occurred (i.e., an approach in swaggering gate with lip-smacking and the come-hither face would not be considered a greeting, unlike in other greeting studies), finding that greetings occurred in about 7% of close proximity approaches (calculated from supplementary material [10]). The comparable rate for the Gorongosa baboons is 14%. The observed percentage of Moremi Game Reserve male-male encounters involving contact falls within our calculated 95% confidence interval (4% to 23%).

Importantly, chacma baboon male-male greeting behaviour appears to align broadly with that of the other COKY baboons. This could mean that (a) their lack of coalition formation is a derived characteristic and that while greetings may function to test cooperative potential in other species they serve a different function in chacma baboons or are a vestigial behaviour, (b) that both coalitionary and greeting behaviour is present to some degree in chacma baboons but has been underestimated and understudied, (c) that greetings reflect a different aspect of relationship quality which may in turn be correlated with cooperation in some but not all *Papio* species, or (c) that the function of greeting behaviour has diverged across the *Papio* genus, but that a base level of ritualized greeting is present across the genus and is likely ancestral. Gorongosa falls in a possible hybridization zone between yellow baboons and northern chacma baboons, so we may expect to see a cline in behaviour similar to the observed cline in morphology [59]. Further systematic study of chacma baboon troops at different distances from the hybridization zone would identify potential effects of hybridization. Gorongosa National Park provides the ideal study site for such work, with 200 troops spread across 3770 km^2^. Our study’s sample size prevents further in-depth comparison with other *Papio* species but does suggest that further research on greeting in southern clade baboons, and particularly in chacma baboons, is warranted.

### 4.2 Signal use and combinations

Through community detection, we identified four clusters of signals that occur together more than expected. The first includes short glances towards the recipient and passing by the recipient without contact or interaction. The second includes observing the recipient, presenting, non-aggressive contact, and arriving and interacting with the recipient upon approach. The third cluster includes observing the recipient’s infant, having physical contact with the infant, and grunting. Cluster four consists of lip-smacking and ear-flattening. Clusters two and three, and three and four, are connected, with parts of this larger connected network appearing very similar to clusters identified when analysing female-female events separately, suggesting that these clusters may be highly related to female-female approaches involving infant contact. The cluster of presenting, interacting upon arrival, and observing during the approach is likely driven by the female-male interactions as it is also detected in this data subset.

The clustering of grunting, infant observation, and infant contact replicates findings of grunting studies across several baboon species, where approachers are found to grunt more when a recipient’s infant is present, possibly as a way of signalling “benign intent” [6,7,51,53]. The juxtaposition of the use of prolonged gaze during approaches that result in interaction (cluster 2) versus the use of short glances during approaches that result in passing by without interaction (cluster 1) suggest that continued observation of the recipient is a potential indicator of intention to interact directly or may be a by-product of the approacher spending time assessing the recipient and context. Cluster 1 (glance toward and pass without contact) appears to be largely driven by female-female, female-male, and male-female interactions interactions, and cluster 2 (arrive and observe during approach) by male-male, male-female and female-male interactions. It is possible that cluster 1 was not identified in male-male interactions due to the limited sample size, and that cluster 2 was masked in female-female interactions by the high proportion of infant-centered interactions. Gaze direction and length are likely associated with the outcome of interactions across all sex combinations. Even if direct gaze does not serve as an intentional signal, primates are generally adept at identifying when they are being looked at, suggesting that direct gaze of an approacher will always serve to transmit information, even if unintentionally [73–75]. The lack of further stereotypical combinations indicates there is significant flexibility in how signals are combined and that approach behaviour in chacma baboons likely cannot be considered “ritualized” [29].

#### 4.2.1 Ear-flattening and lip-smacking

The fourth cluster - lip-smacking and ear-flattening – appears to be driven by both female-female and male-male encounters, according to the observed probability of this signal combination in these sex combinations compared to other sex combinations. The “come hither” or “NEEF” face, which consists of ear-flattening and scalp retraction, is frequently referred to as an affiliative signal in the baboon literature [8,46,76]. When comparing between male and female approachers more generally, rather than splitting further by recipient sex, we find females have a higher probability of performing lip-smacking and males of performing ear-flattening. This indicates that signalling is very clearly affected not only by the sex of the approacher, but also by the sex of the recipient. The combination of lip-smacking and ear-flattening together may be a particularly strong signal of benign intent (male-male encounters can be particularly risky and female-female interactions often involve attempted infant contact), used when a particular outcome, for example physical contact, is desired. The clustering of the lip-smacking and ear-flattening combination together with non-aggressive contact in female-female interactions supports this interpretation. Ear-flattening may be more easily identifiable when observing male than female approachers, but the differences in usage of lips-smacking in male-male approaches and female-female approaches versus all sex combinations could not be explained by this.

Lip-smacking is exhibited by multiple primate species and has been found to be positively associated with affiliative behaviours [54,77,78]. It is one of the most common gestures observed in baboons and is used across a wide variety of contexts [79]. How lip-smacking is combined with other signals influences the outcome of following interactions. In crested macaques, contact after lip-smacking was found to vary based on the signals lip-smacking was combined with, though ear flattening had neither a positive nor negative impact [54]. In chimpanzees, grooming solicitations accompanied by lip-smacking resulted in longer grooming bouts with higher probabilities of reciprocity [78]. Our results indicate that the signals lip-smacking is combined with may also play an important role in baboons and lays the ideal groundwork for further work investigating the outcomes and potential goals of the identified signal clusters.

Similar studies conducted across multiple baboon populations would help determine whether sex-based differences in signal use and combination are consistent within and between species. Using a network-based approach allows for a deeper understanding of the co-occurrence of signals, helping us identify combinations used in specific approach contexts. This approach provides further insight into how combining signals may modify meaning beyond the simple sum of the signals’ individual meanings.

### 4.3 Conclusions

Our results suggest that chacma baboon greeting behaviour aligns with that of the other COKY baboons. Males show some contact behaviour during approaches towards other males, along with many of the other reported greeting behaviours, but it is relatively rare in comparison to Guinea and hamadryas baboons. Within chacma baboons, our comparison of male versus female approaches suggests ear-flattening is used more frequently by males, lip-smacking and presenting by females, and the combination of ear-flattening with lip-smacking is particularly common in male-male and female-female encounters in comparison to other sex combinations. Community detection identified several clusters of signals that co-occur and provides insight into which sex combination approaches are driving the presence of each cluster. The connection between lip-smacking and ear-flattening appears to be particularly relevant to encounters where signalling benign intent may be especially necessary, providing a promising direction for future research.

It is time for a widening of the study of baboon greetings, expanding past the traditional focus on male-male encounters and considering approach behaviour more broadly. Including all proximity events, rather than just instances where individuals “greeted” provides a more encompassing view of approach behaviour and prevents omission of potentially useful information. This approach will allow for more fine-tuned testing of the functions of approach behaviours; knowing in which contexts interactions do not happen during an approach may be just as valuable as knowing in which cases they do. This study provides an example of how widening the methodological framework and using alternative analytical methods, for example network analysis, can give us new insight into the specificity, context, and function of individual behaviours and allow us to identify behaviour clusters used together in a robust way.

## Supporting information

R analysis script

## 5 Acknowledgements

The authors would like to thank collaborator Lucy Baehren, who permitted use of video footage she recorded in Gorongosa National Park, and the staff of Gorongosa National Park for making this research possible, especially Marc Stalmans, Jason Denlinger, and all rangers who worked with us in the field, including Ernesto Xavier and Sérgio Joao Amaral. The authors are most grateful for the vision of Greg Carr and support of the Gorongosa Restoration Project. This work was supported by grant funding from the Boise Trust Fund administered by the Oxford University Department of Zoology and the Owen Aldis Scholarship awarded by the International Society for Human Ethology. Further financial support for fieldwork was provided by St John’s College, University of Oxford and by the Clarendon Fund, University of Oxford.

## Bibliography

1. Dal Pesco F, Fischer J. 2018 Greetings in male Guinea baboons and the function of rituals in complex social groups. Journal of Human Evolution 125, 87–98. (doi:10.1016/j.jhevol.2018.10.007)

2. Dal Pesco F, Fischer J. 2020 On the evolution of baboon greeting rituals. Philosophical Transactions of the Royal Society B 375. (doi:10.1098/rstb.2019.0420)

3. Colmenares F. 1990 Greeting behaviour in male baboons, I: Communication, reciprocity and symmetry. Behaviour 113, 81–116.

4. Mondada L, Meguerditchian A. 2022 Sequence organization and embodied mutual orientations: Openings of social interactions between baboons. Philosophical Transactions of the Royal Society B: Biological Sciences 377, 20210101. (doi:10.1098/rstb.2021.0101)

5. Cheney DL, Seyfarth RM, Silk JB. 1995 The role of grunts in reconciling opponents and facilitating interactions among adult female baboons. Animal Behaviour 50, 249–257.

6. Silk JB, Seyfarth RM, Cheney DL. 2016 Strategic use of affiliative vocalizations by wild female baboons. PLOS ONE 11. (doi:10.1371/journal.pone.0163978)

7. Rendall D, Seyfarth RM, Cheney DL, Owren MJ. 1999 The meaning and function of grunt variants in baboons. Animal Behaviour 57, 583–592.

8. Smuts BB. 1985 Sex and friendship in baboons. London: Transaction Publishers.

9. Hausfater G, Takacs D. 1987 Structure and function of hindquarter presentations in yellow baboons (*Papio cynocephalus*). Ethology 74, 297–319. (doi:10.1111/j.1439-0310.1987.tb00941.x)

10. Kalbitzer U, Heistermann M, Cheney DL, Seyfarth RM, Fischer J. 2015 Social behavior and patterns of testosterone and glucocorticoid levels differ between male chacma and Guinea baboons. Hormones and Behavior 75, 100–110. (doi:10.1016/j.yhbeh.2015.08.013)

11. Fedurek P, Neumann C, Bouquet Y, Mercier S, Magris M, Quintero F, Zuberbühler K. 2019 Behavioural patterns of vocal greeting production in four primate species. Royal Society Open Science 6. (doi:10.1098/rsos.182181)

12. Rodrigues ED, Santos AJ, Hayashi M, Matsuzawa T, Hobaiter C. 2022 Exploring greetings and leave-takings: Communication during arrivals and departures by chimpanzees of the Bossou community, Guinea. Primates 63, 443–461. (doi:10.1007/s10329-021-00957-z)

13. Corewyn LC, Setchell JM. 2019 Greeting behaviors in male *Alouatta palliata* at La Pacifica, Costa Rica. International Journal of Primatology 40, 630–646. (doi:10.1007/s10764-019-00109-7)

14. De Marco A, Cozzolino R, Dessì-Fulgheri F, Thierry B. 2011 Collective arousal when reuniting after temporary separation in Tonkean macaques. American Journal of Physical Anthropology 146, 457–464. (doi:10.1002/ajpa.21606)

15. Dias PAD, Luna ER, Espinosa DC. 2008 The functions of the “greeting ceremony” among male mantled howlers (*Alouatta palliata*) on Agaltepec Island, Mexico. American Journal of Primatology 70, 621–628. (doi:10.1002/ajp.20535)

16. Heesen R et al. 2021 Assessing joint commitment as a process in great apes. iScience 24. (doi:10.1016/J.ISCI.2021.102872)

17. Kutsukake N, Suetsugu N, Hasegawa T. 2006 Pattern, distribution, and function of greeting behavior among black-and-white colobus. International Journal of Primatology 27. (doi:10.1007/s10764-006-9072-x)

18. Laporte MNC, Zuberbühler K. 2010 Vocal greeting behaviour in wild chimpanzee females. Animal Behaviour 80, 467–473. (doi:10.1016/J.ANBEHAV.2010.06.005)

19. Luef EM, Pika S. 2017 Reciprocal greeting in chimpanzees (*Pan troglodytes*) at the ngogo community. Journal of Neurolinguistics 43, 263–273. (doi:10.1016/J.JNEUROLING.2016.11.002)

20. Luef EM, Pika ·S. 2019 Social relationships and greetings in wild chimpanzees (*Pan troglodytes*): Use of signal combinations. Primates 60, 507–515. (doi:10.1007/s10329-019-00758-5)

21. Okamoto K, Agetsuma N, Kojima S. 2001 Greeting behavior during party encounters in captive chimpanzees. Primates 42, 161–165. (doi:10.1007/BF02558143)

22. Nakamura M. 2022 Greetings among female chimpanzees in Mahale, Tanzania. American Journal of Primatology **n/a**, e23417. (doi:10.1002/ajp.23417)

23. Schaffner CM, Aureli F. 2005 Embraces and grooming in captive spider monkeys. International Journal of Primatology 26. (doi:10.1007/s10764-005-6460-6)

24. Scheumann M, Linn S, Zimmermann E. 2017 Vocal greeting during mother-infant reunions in a nocturnal primate, the gray mouse lemur (*Microcebus murinus*). Scientific Reports 7, 1–7. (doi:10.1038/s41598-017-10417-8)

25. de Waal FBM, Roosmalen A van. 1979 Reconciliation and consolation among chimpanzees. Behavioral Ecology and Sociobiology 5, 55–66. (doi:10.1007/BF00302695)

26. Kavanagh E et al. 2021 Dominance style is a key predictor of vocal use and evolution across nonhuman primates. Royal Society Open Science 8, 210873. (doi:10.1098/rsos.210873)

27. Whitham JC, Maestripieri D. 2003 Primate rituals: The function of greetings between male Guinea baboons. Ethology 109, 847–859. (doi:10.1046/j.0179-1613.2003.00922.x)

28. Maestripieri D. 2005 Gestural communication in three species of macaques (*Macaca mulatta, M. nemestrina, M. arctoides*): Use of signals in relation to dominance and social context. Gesture 5, 57–73.

29. Rossano MJ. 2012 The essential role of ritual in the transmission and reinforcement of social norms. Psychological Bulletin 138, 529–549. (doi:10.1037/a0027038)

30. Whitehouse H, Lanman JA. 2014 The ties that bind us: Ritual, fusion, and identification. Current Anthropology 55. (doi:10.1086/678698)

31. Rossano MJ. 2015 The evolutionary emergence of costly rituals. PaleoAnthropology, 78–100. (doi:10.4207/PA.2015.ART97)

32. Bobe R, Coelho J, Carvalho S, Leakey M. 2022 Early hominins and paleoecology of the Koobi Fora Formation, Lake Turkana Basin, Kenya. In African paleoecology and human evolution (ed SC Reynolds), pp. 311–331. Cambridge: Cambridge University Press.

33. Roos C, Knauf S, Chuma IS, Maille A, Callou C, Sabin R, Portela Miguez R, Zinner D. 2021 New mitogenomic lineages in *Papio* baboons and their phylogeographic implications. American Journal of Physical Anthropology 174, 407–417. (doi:10.1002/AJPA.24186)

34. Zinner D, Wertheimer J, Liedigk R, Groeneveld LF, Roos C. 2013 Baboon phylogeny as inferred from complete mitochondrial genomes. American Journal of Physical Anthropology 150, 133–140. (doi:10.1002/ajpa.22185)

35. Ey E, Rahn C, Hammerschmidt K, Fischer J. 2009 Wild female olive baboons adapt their grunt vocalizations to environmental conditions. Ethology 115, 493–503.

36. Graham KE, Badihi G, Safryghin A, Grund C, Hobaiter C. 2022 A socio-ecological perspective on the gestural communication of great ape species, individuals, and social units. Ethology Ecology & Evolution 34, 235–259. (doi:10.1080/03949370.2021.1988722)

37. Jolly CJ, Phillips-Conroy JE. 2006 Testicular size, developmental trajectories, and male life history strategies in four baboon taxa. In Reproduction and fitness in baboons: Behavioral, ecological, and life history perspectives (eds L Swedell, SR Leigh), pp. 257–275. Springer.

38. Plavcan JM, Ruff CB. 2008 Canine size, shape, and bending strength in primates and carnivores. 84, 65–84. (doi:10.1002/ajpa.20779)

39. Thorén S, Lindenfors P, Kappeler PM. 2006 Phylogenetic analyses of dimorphism in primates: Evidence for stronger selection on canine size than on body size. American Journal of Physical Anthropology 130, 50–59. (doi:10.1002/ajpa.20321)

40. Henzi SP, Barrett L. 2003 Evolutionary ecology, sexual conflict, and behavioral differentiation among baboon populations. Evolutionary Anthropology 12, 217–230. (doi:10.1002/evan.10121)

41. Henzi SP, Barrett L. 2005 The historical socioecology of savanna baboons (*Papio hamadryas*). Journal of Zoology 265, 215–226. (doi:10.1017/S0952836904006399)

42. Saayman GS. 1971 Behaviour of the adult males in a troop of free-ranging chacma baboons (*Papio ursinus*). Folia Primatologica 15, 36–57.

43. Fischer J et al. 2017 Charting the neglected West: The social system of Guinea baboons. (doi:10.1002/ajpa.23144)

44. Kummer H. 1984 From laboratory to desert and back: A social system of hamadryas baboons. Animal Behaviour 32, 965–971. (doi:10.1016/S0003-3472(84)80208-0)

45. Patzelt A, Kopp GH, Ndao I, Kalbitzer U, Zinner D, Fischer J. 2014 Male tolerance and male male bonds in a multilevel primate society. Proceedings of the National Academy of Sciences 111, 14740–14745. (doi:10.1073/pnas.1405811111)

46. Smuts BB, Watanabe JM. 1990 Social relationships and ritualized greetings in adult male baboons (*Papio cynocephalus anubis*). International Journal of Primatology 11, 147–.

47. Fraser Ó, Plowman AB. 2007 Function of notification in *Papio hamadryas*. International Journal of Primatology 28, 1439–1448. (doi:10.1007/s10764-007-9185-x)

48. Colmenares F. 1991 Greeting, aggression, and coalitions between male baboons: Demographic correlates. Primates 32, 453. (doi:10.1007/BF02381936)

49. Henzi SP, Clarke PMR, Barrett L, Noë R, Jolly CJ. 2008 Cooperation and coalition formation in humans and primates: The genetics and biogeography of coalition formation in savanna baboons.

50. Hall KRL. 1962 The sexual, agonistic and derived social behaviour patterns of the wild chacma baboon, *Papio Ursinus*. Proceedings of the Zoological Society of London 139, 283–327. (doi:10.1111/j.1469-7998.1962.tb01831.x)

51. Silk JB, Roberts ER, Städele V, Strum SC. 2018 To grunt or not to grunt: Factors governing call production in female olive baboons, *Papio anubis*. PLOS ONE 13. (doi:10.1371/journal.pone.0204601)

52. Cheney DL, Seyfarth RM. 1997 Reconciliatory grunts by dominant female baboons influence victims’ behaviour. Animal Behaviour 54, 409–418. (doi:10.1006/anbe.1996.0438)

53. Faraut L, Siviter H, Dal Pesco F, Fischer J. 2019 How life in a tolerant society affects the usage of grunts: Evidence from female and male Guinea baboons. Animal Behaviour 153, 83–93. (doi:10.1016/j.anbehav.2019.05.003)

54. Micheletta J, Engelhardt A, Matthews L, Agil M, Waller BM. 2013 Multicomponent and multimodal lipsmacking in crested macaques (*Macaca nigra*). American Journal of Primatology 75, 763–773. (doi:10.1002/ajp.22105)

55. Mielke A, Waller BM, Pérez C, Rincon AV, Duboscq J, Micheletta J. 2021 NetFACS: Using network science to understand facial communication systems. Behavior Research Methods (doi:10.3758/s13428-021-01692-5)

56. Aychet J, Blois-Heulin C, Lemasson A. 2021 Sequential and network analyses to describe multiple signal use in captive mangabeys. Animal Behaviour 182, 203–226. (doi:10.1016/j.anbehav.2021.09.005)

57. Bobe R et al. 2021 The first Miocene fossils from coastal woodlands in the southern East African Rift. bioRxiv (doi:10.1101/2021.12.16.472914)

58. Macgregor D. 2015 History of the development of the East African Rift System: A series of interpreted maps through time. Journal of African Earth Sciences 101, 232–252. (doi:10.1016/j.jafrearsci.2014.09.016)

59. Martinez FI et al. 2019 A missing piece of the *Papio* puzzle: Gorongosa baboon phenostructure and intrageneric relationships. Journal of Human Evolution 130, 1–20. (doi:10.1016/j.jhevol.2019.01.007)

60. Tinley KL. 1977 Framework of the Gorongosa ecosystem. PhD thesis.

61. Habermann JM et al. 2018 Gorongosa by the sea: First Miocene fossil sites from the Urema Rift, central Mozambique, and their coastal paleoenvironmental and paleoecological contexts. Palaeogeography, Palaeoclimatology, Palaeoecology

62. Stalmans M, Beilfuss R. 2008 Landscapes of the Gorongosa National Park.

63. Hammond P, Lewis-Bevan L, Biro D, Carvalho S. 2022 Risk perception and terrestriality in primates: A quasi-experiment through habituation of chacma baboons (*Papio ursinus*) in Gorongosa National Park, Mozambique. American Journal of Biological Anthropology 179, 48–59. (doi:10.1002/ajpa.24567)

64. Baehren L, Carvalho S. 2022 Yet another non-unique human behaviour: Leave-taking in wild chacma baboons (*Papio ursinus*). Animals 12, 2577. (doi:10.3390/ani12192577)

65. Colmenares F. 1991 Greeting behaviour between male baboons: Oestrous females, rivalry and negotiation. Animal Behaviour 41, 49–60.

66. Silk JB. 2013 CABS Project Research Protocol Guide. Uaso Ngiro Baboon Project.

67. Muschinski J, Carvalho S. 2022 Papio ursinus *approach behaviour ethogram, Gorongosa National Park, Mozambique*. 1st edn. Gorongosa National Park, Sofala, Mozambique: Paleo-Primate Project, Gorongosa National Park. (doi:10.5281/zenodo.7314291)

68. Muschinski J, Carvalho S. 2022 Chacma baboon approach behaviour dataset, Gorongosa National Park, Mozambique. (doi:10.5281/zenodo.7330446)

69. Friard O, Gamba M. 2016 BORIS: A free, versatile open-source event-logging software for video/audio coding and live observations. Methods in Ecology and Evolution 7, 1325–1330. (doi:10.1111/2041-210X.12584)

70. Silge J, Robinson D. 2017 Text Mining with R: A Tidy Approach. O’Reilly Media. See https://www.tidytextmining.com/.

71. Canty A, Ripley B. 2016 Package ‘boot’. CRAN. See https://cran.microsoft.com/snapshot/2016-04-15/web/packages/boot/boot.pdf.

72. Newman MEJ. 2004 Fast algorithm for detecting community structure in networks. Physical Review E 69, 066133. (doi:10.1103/PhysRevE.69.066133)

73. Bourjade M, Meguerditchian A, Maille A, Gaunet F, Vauclair J. 2014 Olive baboons, *Papio anubis*, adjust their visual and auditory intentional gestures to the visual attention of others. Animal Behaviour 87, 121–128. (doi:10.1016/J.ANBEHAV.2013.10.019)

74. Emery NJ. 2000 The eyes have it: The neuroethology, function and evolution of social gaze. Neuroscience and Biobehavioral Reviews 24, 581–604.

75. Harrod EG, Coe CL, Niedenthal PM. 2020 Social structure predicts eye contact tolerance in nonhuman primates: Evidence from a crowd-sourcing approach. Scientific Reports, 1–9. (doi:10.1038/s41598-020-63884-x)

76. Smuts BB. 2002 Gestural communication in olive baboons and domestic dogs. In The cognitive animal (eds M Bekoff, C Allen, GM Burghardt), pp. 301–306. London: The MIT Press.

77. Easley SP, Coelho AM. 1991 Is lipsmacking an indicator of social status in baboons? Folia Primatologica 56, 190–201. (doi:10.1159/000156547)

78. Fedurek P, Slocombe KE, Hartel JA, Zuberbühler K. 2015 Chimpanzee lip-smacking facilitates cooperative behaviour. Scientific Reports 5, 13460. (doi:10.1038/srep13460)

79. Molesti S, Meguerditchian A, Bourjade M. 2020 Gestural communication in olive baboons (*Papio anubis*): repertoire and intentionality. Animal Cognition 23, 19–40. (doi:10.1007/s10071-019-01312-y)

80. Rogers J et al. 2019 The comparative genomics and complex population history of Papio baboons. Science Advances 5.

81. Petersdorf M, Weyher AH, Kamilar JM, Dubuc C, Higham JP. 2019 Sexual selection in the Kinda baboon. Journal of Human Evolution 135, 102635. (doi:10.1016/j.jhevol.2019.06.006)

82. Swedell L, Saunders J, Schreier A, Davis B, Tesfaye T, Pines M. 2011 Female “dispersal” in hamadryas baboons: Transfer among social units in a multilevel society. American Journal of Physical Anthropology 145, 360–370. (doi:10.1002/ajpa.21504)

83. Henzi SP, Weingrlll T, Barretta L. 1999 Male behaviour and the evolutionary ecology of chacma baboons. South African Journal of Science 95, 240–242.

84. Pelaez F. 1982 Greeting movements among adult males in a colony of baboons: *Papio hamadryas, P. cynocephalus* and their hybrids. Primates 23, 233–244.

