## Supplemental tables and figures for "A network-based analysis of signal use during approach interactions across sexes in chacma baboons (*Papio ursinus griseipes*)"

### Supplementary Materials - Results

#### Community Detection [Section 3.2]:

##### Full Ethogram Community Detection:

Three clusters were identified using the expanded ethogram, with an overall modularity of 0.61 (see figure S.1). The clusters were (1) passing without contact paired with glancing toward the recipient during the approach, (2) observing an infant, grunting, ear-flattening, and lip-smacking, and (3) arriving (classified as an approach that ends in the individual stopping at and/or interacting with the recipient rather than diverting or passing without contact), observing the recipient during the approach, presenting, and attempting to grab the recipient's infant.

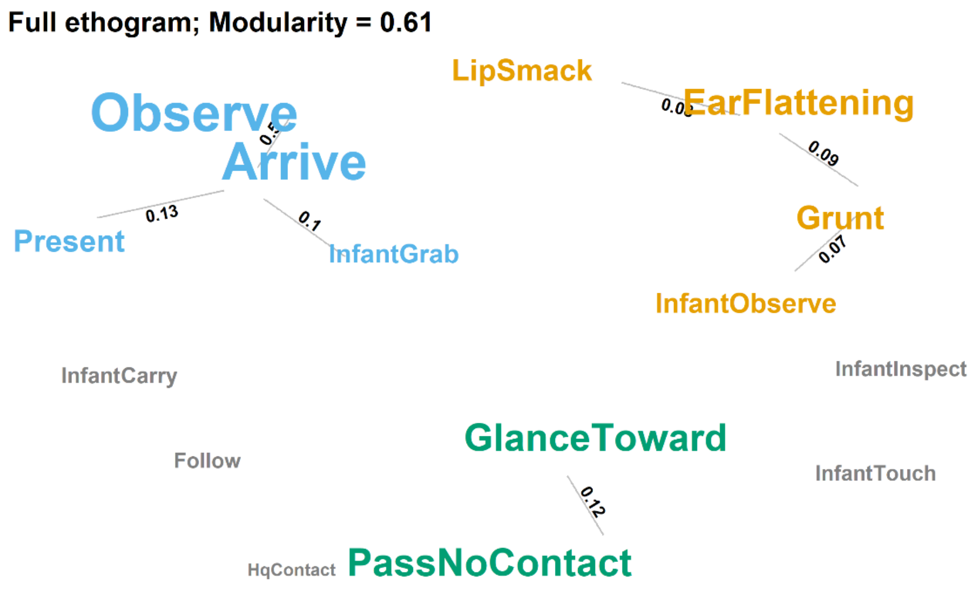


*Figure S.1: Community detection across all greetings with the full ethogram. Linked and coloured behaviours are detected clusters, with edges labelled with the combination's observed probability*

##### Sex Combination Subset Community Detection:

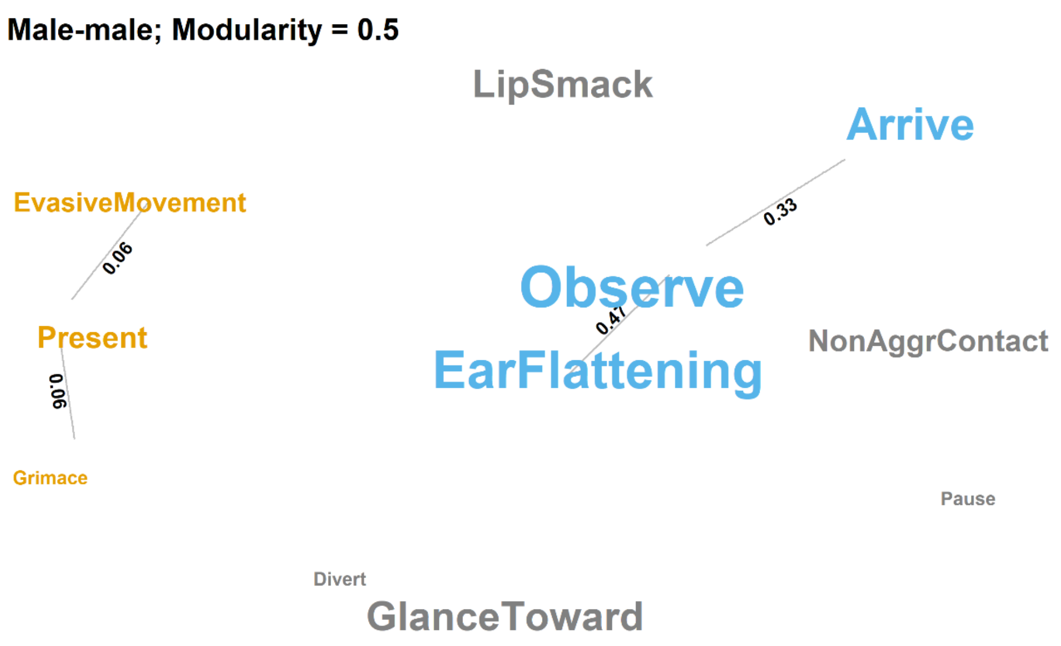


*Figure S.2: Community detection across male-male approaches with the collapsed ethogram. Linked and coloured behaviours are detected clusters, with edges labelled with the combination's observed probability*

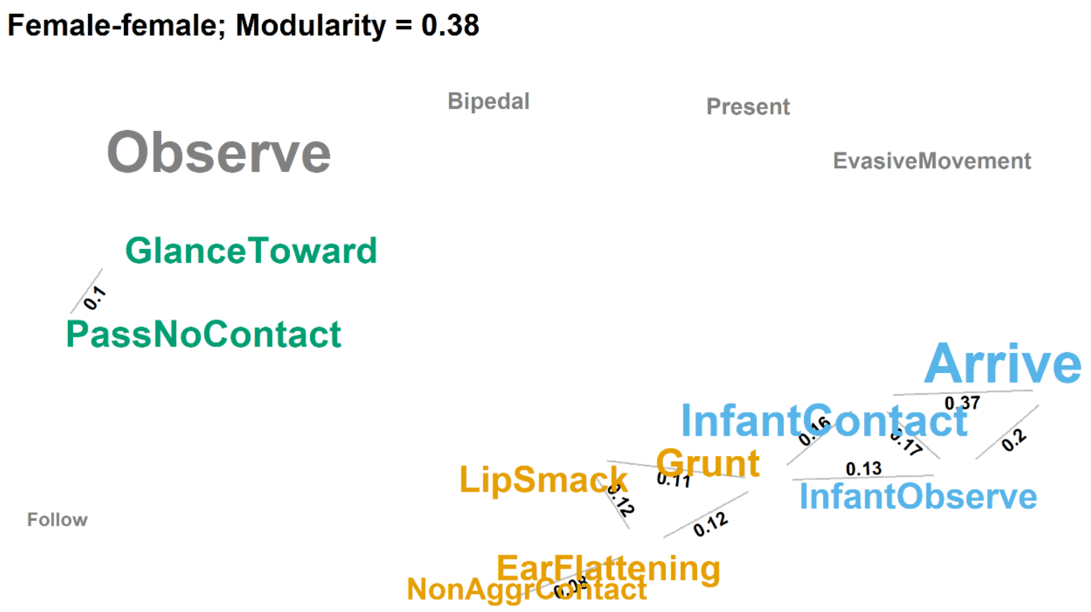


*Figure S.3: Community detection across female-female approaches with a collapsed ethogram. Linked and coloured behaviours are detected clusters, with edges labelled with the combination's observed probability*

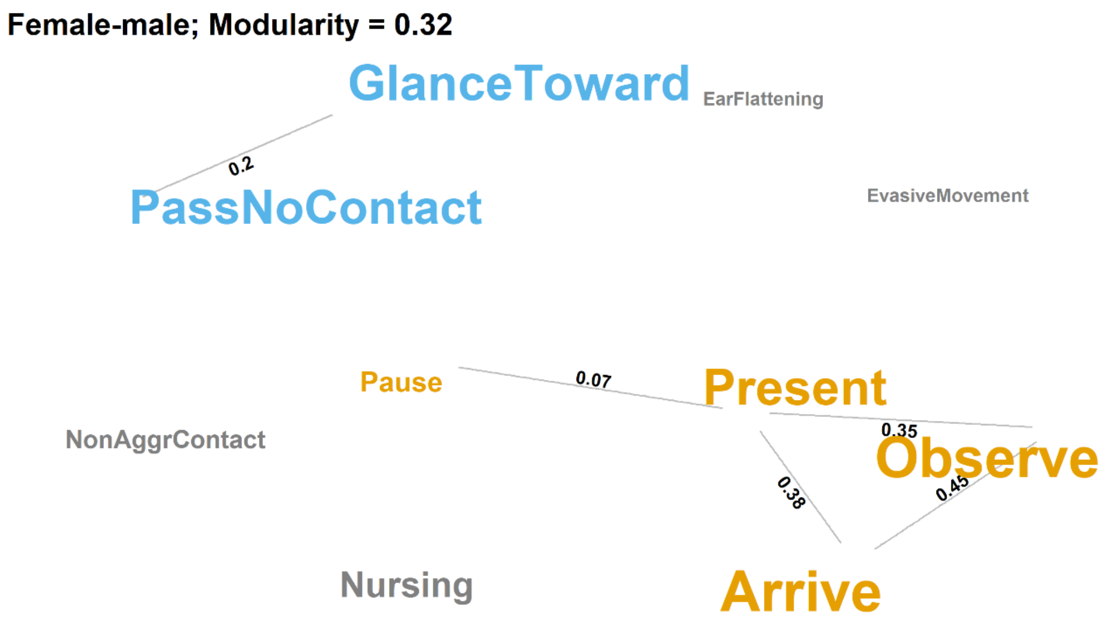


*Figure S.4: Community detection across female-male approaches with a collapsed ethogram. Linked and coloured behaviours are detected clusters, with edges labelled with the combination's observed probability*

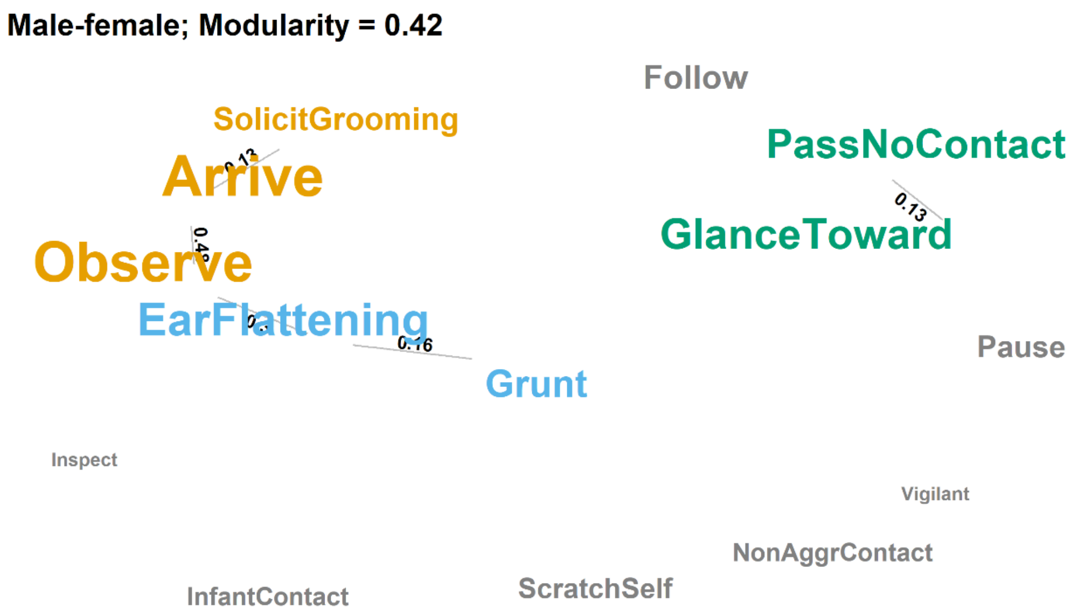


*Figure S.5: Community detection across male-female approaches with a collapsed ethogram. Linked and coloured behaviours are detected clusters, with edges labelled with the combination's observed probability*

#### Signal Use Across Sex Combinations [Section 3.3.2]:

Table S.1: Signal use in female-female approaches compared to a bootstrapped sample of approaches in all sex combinations

| Combination | Combination size | Count | Expected probability | Observed probability | Effect size | p-value | Probability increase |
|---|---|---|---|---|---|---|---|
| **LipSmack** | 1 | 36 | 0.063 | 0.217 | 0.154 | 0 | 3.435 |
| Arrive_LipSmack | 2 | 33 | 0.036 | 0.199 | 0.163 | 0 | 5.587 |
| LipSmack_Observe | 2 | 29 | 0.058 | 0.175 | 0.116 | 0 | 2.987 |
| EarFlattening_Grunt | 2 | 20 | 0.071 | 0.12 | 0.049 | 0 | 1.693 |
| EarFlattening_LipSmack | 2 | 20 | 0.05 | 0.12 | 0.07 | 0 | 2.408 |
| Grunt_LipSmack | 2 | 19 | 0.009 | 0.114 | 0.105 | 0 | 12.623 |
| EarFlattening_InfantObserve | 2 | 11 | 0.009 | 0.066 | 0.057 | 0 | 7.215 |
| EarFlattening_InfantInspect | 2 | 10 | 0.004 | 0.06 | 0.056 | 0 | 14.399 |
| EarFlattening_InfantCarry | 2 | 9 | 0.004 | 0.054 | 0.05 | 0 | 12.959 |
| EarFlattening_InfantGrab | 2 | 9 | 0.018 | 0.054 | 0.036 | 0 | 3 |
| GlanceToward_LipSmack | 2 | 6 | 0.013 | 0.036 | 0.023 | 0 | 2.758 |
| LipSmack_Present | 2 | 6 | 0.014 | 0.036 | 0.022 | 0.001 | 2.59 |
| EarFlattening_InfantTouch | 2 | 5 | 0.009 | 0.03 | 0.021 | 0 | 3.279 |
| Grunt_HqContact | 2 | 5 | 0.004 | 0.03 | 0.026 | 0 | 7.456 |
| LipSmack_HqContact | 2 | 4 | 0.004 | 0.024 | 0.02 | 0 | 5.965 |
| Arrive_LipSmack_Observe | 3 | 26 | 0.031 | 0.157 | 0.126 | 0 | 5.065 |
| Arrive_EarFlattening_Grunt | 3 | 18 | 0.045 | 0.108 | 0.064 | 0 | 2.432 |
| Arrive_EarFlattening_LipSmack | 3 | 18 | 0.032 | 0.108 | 0.077 | 0 | 3.438 |
| EarFlattening_Grunt_Observe | 3 | 15 | 0.062 | 0.09 | 0.028 | 0.032 | 1.451 |
| EarFlattening_LipSmack_Observe | 3 | 15 | 0.045 | 0.09 | 0.045 | 0.002 | 1.992 |

| | | | | | | | |
|---|---|---|---|---|---|---|---|
| Grunt_LipSmack_Observe | 3 | 14 | 0.009 | 0.084 | 0.075 | 0 | 9.301 |
| Arrive_EarFlattening_InfantGrab | 3 | 9 | 0.018 | 0.054 | 0.036 | 0 | 3 |
| EarFlattening_Grunt_InfantObserve | 3 | 9 | 0.009 | 0.054 | 0.045 | 0 | 5.903 |
| EarFlattening_InfantObserve_Observe | 3 | 8 | 0.009 | 0.048 | 0.039 | 0 | 5.247 |
| Arrive_EarFlattening_HqContact | 3 | 7 | 0.021 | 0.042 | 0.021 | 0.009 | 1.994 |
| EarFlattening_Grunt_InfantGrab | 3 | 7 | 0.018 | 0.042 | 0.024 | 0 | 2.333 |
| Arrive_GlanceToward_LipSmack | 3 | 6 | 0.004 | 0.036 | 0.032 | 0 | 8.948 |
| Arrive_LipSmack_Present | 3 | 6 | 0.014 | 0.036 | 0.022 | 0.001 | 2.59 |
| EarFlattening_InfantGrab_Observe | 3 | 6 | 0.018 | 0.036 | 0.018 | 0.002 | 2 |
| EarFlattening_Grunt_HqContact | 3 | 5 | 0.004 | 0.03 | 0.026 | 0 | 7.456 |
| GlanceToward_LipSmack_Observe | 3 | 5 | 0.013 | 0.03 | 0.017 | 0 | 2.298 |
| Grunt_Observe_HqContact | 3 | 5 | 0.004 | 0.03 | 0.026 | 0 | 7.456 |
| LipSmack_Observe_Present | 3 | 5 | 0.014 | 0.03 | 0.016 | 0.004 | 2.158 |
| Arrive_EarFlattening_Follow | 3 | 4 | 0.004 | 0.024 | 0.02 | 0 | 5.965 |
| Arrive_Grunt_HqContact | 3 | 4 | 0.004 | 0.024 | 0.02 | 0 | 5.965 |
| EarFlattening_InfantInspect_InfantCarry | 3 | 4 | 0.004 | 0.024 | 0.02 | 0 | 5.76 |
| EarFlattening_InfantObserve_InfantTouch | 3 | 4 | 0.009 | 0.024 | 0.015 | 0 | 2.623 |
| GlanceToward_Grunt_LipSmack | 3 | 4 | 0.009 | 0.024 | 0.015 | 0 | 2.657 |
| LipSmack_Observe_HqContact | 3 | 4 | 0.004 | 0.024 | 0.02 | 0 | 5.965 |
