## Supplementary material for "A network-based analysis of signal use during approach interactions across sexes in chacma baboons (*Papio ursinus griseipes*)": R analysis script: Muschinski_et_al_analysis_script.R

```
# File:     Analysis script
# Project:  A network-based analysis of signal use during approach interactions across sexes in chacma baboons (Papio ursinus)

# INSTALL AND LOAD PACKAGES ################################

#install.packages("pacman")
# pacman must already be installed; then load contributed
# packages (including pacman) with pacman

set.seed(233)


#load packages
pacman::p_load(pacman, patchwork, NetFACS, tidyverse, magrittr, ggplot2, dplyr, 
               grid, gridExtra, igraph, ggraph, grDevices, tidygraph, tinytex, 
               haven, sjstats, ROCR, performance, patchwork, flextable, boot, data.table, RColorBrewer)

packageVersion('NetFACS') # should be 0.5.0


#set filepath for newest export of data
filepath_max = "Paper_1/processed-data/NetFACS_DF_max_cats_21.01.22_export_no-unassigned.csv"
filepath_min = "Paper_1/processed-data/NetFACS_DF_min_cats_21.01.22_export_no-unassigned.csv"

filepath_video_info = "Paper_1/processed-data/video_info_21.01.22_export_no-unassigned.csv"

# LOAD DATA ####################################

max_DF <- read.csv(filepath_max) %>%
  subset(, -c(1)) #drop observation ID column

min_DF <- read.csv(filepath_min) %>%
  subset(, -c(1)) #drop obs id column

video_info <- read.csv(filepath_video_info, header = T, na.strings="") %>%
  mutate(Greeting_type = paste(Greeter_sex, Greetee_sex)) # create a column that has sex combination of greeting

## View data #################################################

head(max_DF)

head(min_DF)

head(video_info)


# PREPARE DATA #####################################################

# We remove behaviours that have 4 or fewer occurrences, because not meaningful interpretation is available for them
max_DF <- max_DF[, colSums(max_DF) > 4]  #present in at least 1% of interactions
min_DF <- min_DF[, colSums(min_DF) > 4]

head(max_DF)


#Underscores not allowed to be in variable names, so renaming variables to CamelCase


print(names(max_DF))

print(names(min_DF))


colnames(max_DF) <- c("Arrive", "Bipedal", "Chase", "Divert", "DodgeFlinch", 
                      "EarFlattening", "Follow", "Forage", "GlanceToward", 
                      "Grimace", "Grunt", "InfantGrab", "InfantInspect", 
                      "InfantObserve", "InfantTouch", "Inspect", "LipSmack", 
                      "LookAway", "Lunge", "Mount", "Nursing", "Observe", 
                      "PassNoContact", "Pause", "ScratchSelf", "Sniff", 
                      "SolicitGrooming", "Vigilant", "EvasiveBodyShift", "HqContact",
                      "TouchNStroke", "Embrace", "Present", "Bump", "InfantCarry")

print(names(max_DF)) #check


colnames(min_DF) <- c("Arrive", "Bipedal", "Divert", "EarFlattening", "Follow",
                      "Forage", "GlanceToward", "Grimace", "Grunt", "InfantObserve",
                      "Inspect", "LipSmack", "Nursing", "Observe", "PassNoContact",
                      "Pause", "ScratchSelf", "Sniff", "SolicitGrooming", "Vigilant",
                      "EvasiveMovement", "ThreatOrAggrContact", "NonAggrContact",
                      "InfantContact", "Present")

print(names(min_DF)) #check


# DESCRIPTIVE STATS ################################################

## basic counts #################
nrow(video_info) #total number that meet face visibility requirements


table(is.na(video_info$Greeter)) #number where approacher is vs is not identified

table(is.na(video_info$Greetee)) #number where recipient is vs is not identified

sum(is.na(video_info$Greeter) & is.na(video_info$Greetee)) #where both are not


table(video_info$Greeter_sex, video_info$Greetee_sex) #count of interactions by sex


## average event size ################

# Mean number of behaviours for each group when leaving out arrive, divert, PassNoContact

glimpse(max_DF)

#create new dataframe for this part since we aren't including these behaviours
avgs_DF <- max_DF%>%
  select(-c("Arrive", "Divert", "Nursing", "PassNoContact", "Forage"))

#create new column that is the sum of how many unique behaviours happened per row
avgs_DF$indsum <- rowSums(avgs_DF[,])

#combine this dataframe with video_info dataframe
avgs_DF <- avgs_DF %>%
  cbind(video_info) 

#table of event size means and sd's by sex combo
ev_size <- avgs_DF %>%
  group_by(Greeting_type) %>% #group by the greeting type (sex combo)
  summarise(
    Mean = round(mean(indsum),2),  #create new Mean variable
    SD = round(sd(indsum), 2) #create new SD variable
    )

rm(avgs_DF, ev_size)


# 3.1 MALE MALE ANALYSIS ##########################################

## import MM dataset ################################
filepath_MM = "Paper_1/processed-data/NetFACS_DF_all_MM_events_greeter_greetee.csv"
MM_DF <- read.csv(filepath_MM)


## Calculate percentages #########################

# percentage of mm greetings having any contact
contactP = round(sum(MM_DF$contact)/sum(MM_DF$greet), 3)*100

#percentage of mm greetings having intense contact
intenseP = round(sum(MM_DF$intense)/sum(MM_DF$greet), 3)*100

# percentage that are reciprocal 
reciprocalP = round(sum(MM_DF$reciprocal)/sum(MM_DF$greet), 3)*100

# percentage of all MM proximity events (not just greetings) that involve contact
contact_all = round(sum(MM_DF$contact)/nrow(MM_DF),2)*100


## Bootstrap confidence intervals ###############


meanfun <- function(data, i){
  d <- data[i]
  return(mean(d))   
}

# create dataframe that only includes those classified as greetings
df.2 <- copy(MM_DF) %>%
  filter(greet == 1)

set.seed(55) #same seed as in RMarkdown file that generates paper
#confidence interval for contact as percentage of all MM interactions
c_all <- boot(MM_DF$contact, statistic=meanfun, R=5000)
c_all_ci <- boot.ci(c_all, conf=0.95, type="norm")


#confidence interval for contact as percentage of only greeting 
c_greet <- boot(df.2$contact, statistic = meanfun, R = 5000)
c_greet_ci <- boot.ci(c_greet, conf = 0.95, type = "norm")


#confidence interval for intense contact as percentage of only greeting
c_intense <- boot(df.2$intense, statistic = meanfun, R = 5000)
c_intense_ci <- boot.ci(c_intense, conf = 0.95, type = "norm")


#confidence interval for reciprocal greeting of only greeting
c_recip <- boot(df.2$reciprocal, statistic = meanfun, R = 5000)
c_recip_ci <- boot.ci(c_recip, conf = 0.95, type = "norm")


rm(contact_all, contactP, intenseP, reciprocalP, c_all, c_all_ci, c_greet,c_greet_ci, c_intense, c_intense_ci, c_recip, c_recip_ci, df.2)


# 3.2 COMMUNITY DETECTION    ################################################

# plotting function 
# based on the netfacs community detection plot function, but slightly modified
# used in creation of the supplementary materials figures


complot <- function(netfacs.graph,
                    title = "network",
                    hide.unconnected = TRUE) {
  g <- netfacs.graph
  nodes <- NULL # to avoid R CMD check when calling activate(nodes)
  
  if (hide.unconnected) {
    g <- 
      g %>% 
      activate(nodes) %>% 
      filter(.data$element.prob != 0 | !is.na(.data$element.prob))
  }
  
  
  net.un <- g
  # other clustering algorithms exist, eg walktrap
  net.community <- cluster_fast_greedy(net.un) 
  # modularity measure. Above 0.3 is good modularity
  modular <- round(modularity(net.community), 2) 
  
  net.com <- data.frame(
    name = net.community$names,
    community = net.community$membership
  )
  
  
  col_tb <- table(net.com['community'])
  
  col_tb<- data.frame(col_tb)
  
  
  colour.group <- function(x) {
    if (col_tb$Freq[x] > 1){
      group = as.integer(x+1)
      return(group)}
    return(as.integer(1))
  }
  
  
  
  net.com$col_group <- purrr::map_chr(net.com$community, colour.group)
  
  cbPalette <- c("#808080", "#E69F00", "#56B4E9", "#009E73", "#F0E442", "#0072B2", "#D55E00", "#CC79A7")
  colornum <- net.com$col_group %>%
    unique() %>%
    length()
  color <- head(cbPalette, colornum)
  shape <- sample(rep(15:19, 10))
  
  p <- 
    g %>% 
    ggraph(layout = "igraph", 
           algorithm = 'kk') +
    geom_edge_link(aes(label = round(.data$observed.prob, 2)),
                   arrow = NULL,
                   colour = "grey",
                   fontface = 'bold',
                   end_cap = circle(5, "mm"),
                   start_cap = circle(5, "mm"),
                   label_dodge = unit(2, "mm"),
                   angle_calc = "along",
                   show.legend = FALSE) +
    geom_node_text(aes(label = .data$name,
                       color = net.com$col_group,
                       size = .data$element.prob,
                       fontface = "bold"),
                   show.legend = FALSE,
                   repel = TRUE) +
    scale_size(range = c(4, 12)) +
    ggtitle(paste(c(title, "; Modularity = ", modular), collapse = "")) +
    theme_graph(base_family = "sans") +
    scale_color_manual(values = color)
  
  return(p)
}


## Max categories  ####################

set.seed(55) #resetting seed for each analysis so they can be run 
# independently and still be reproducible

#run bootstrapping for expanded ethogram
max.all.greet<- netfacs(
  data = max_DF,
  condition = NULL,
  ran.trials = 2000,
  combination.size = 3
)


#do community detection for the expanded ethogram
max.all.net <- netfacs_network(netfacs.data = max.all.greet,
                               significance = 0.05, # cut off value of 0.05 due to low sample size
                               min.count = 17, #happens in at least 5% of interactions
                               link = "unweighted",
                               min.prob = 0.05)


#view the community detection plot
complot(max.all.net, title = "Full ethogram", hide.unconnected = TRUE)

#save the bootstrapped sample of the maximum ethogram dataframe
saveRDS(max.all.greet, "Paper_1/RDS_objects/max_all_greet")

#network summary so it's in a form that can be read as a table
max_net_sum <- network_summary(max.all.net)

#save the network summary
saveRDS(max_net_sum, "Paper_1/RDS_objects/max_all_net")

#save the community detection figure
ggsave("max_net_complot.png", device = "png", path = "Paper_1/figures/", width = 10, height = 6)

#remove unnecessary objects from environment
rm(max.all.greet, max.all.net)


## Min categories ####

set.seed(55) #resetting seed for each analysis so they can be run 
# independently and still be reproducible

#run bootstrapping
min.all.greet<- netfacs(
  data = min_DF,
  condition = NULL,
  ran.trials = 2000,
  combination.size = 3
)

#run community detection
min.all.net <- netfacs_network(netfacs.data = min.all.greet,
                               significance = 0.05, # cut off value of 0.05 due to low sample size
                               min.count = 17, #happens in at least 5% of interactions
                               link = "unweighted",
                               min.prob = 0.05)

# plot it
complot(min.all.net, hide.unconnected = TRUE, title = "Collapsed ethogram")

# save the bootstrapped object
saveRDS(min.all.greet, "Paper_1/RDS_objects/min_all_greet")

# get the network summary
min_net_sum <- network_summary(min.all.net)

# save the network summary
saveRDS(min_net_sum, "Paper_1/RDS_objects/min_all_net")


### export to igraph object #############
# create an edge table
min.all.net.table <- igraph::as_data_frame(min.all.net)

min.all.net.table


# save as CSV file
write.csv(min.all.net.table, "Paper_1/processed-data/min_net_table.csv")

# Community detection plot for paper was made in Gephi using the above saved table


## Supplementary figures #############
# the below are used to create the plots for the supplementary materials

### Male-male ################
set.seed(55)
#to look at only male-male
MM_subset <- min_DF %>%
  cbind(video_info) %>%
  filter(Greeting_type == "male male") %>%
  select(c(1:25))


MM_min_all_greet<- netfacs(
  data = MM_subset,
  condition = NULL,
  ran.trials = 2000,
  combination.size = 3,
  use_parallel = TRUE
)


MM_min_all_net <- netfacs_network(netfacs.data = MM_min_all_greet,
                                  significance = 0.05, # cut off value of 0.05 due to low sample size
                                  min.count = 2, #happens in at least 5% of the 37 observations
                                  link = "unweighted",
                                  min.prob = 0.05)

complot(MM_min_all_net, title = "Male-male")


ggsave("MM_complot.png", device = "png", path = "Paper_1/figures/", width = 10, height = 6)

### Female-female ################

set.seed(55)
#to look at only female-female
FF_subset <- min_DF %>%
  cbind(video_info) %>%
  filter(Greeting_type == "female female") %>%
  select(c(1:25))


FF_min_all_greet<- netfacs(
  data = FF_subset,
  condition = NULL,
  ran.trials = 2000,
  combination.size = 3,
  use_parallel = TRUE
)

FF_min_all_net <- netfacs_network(netfacs.data = FF_min_all_greet,
                                  significance = 0.05, # cut off value of 0.05 due to low sample size
                                  min.count = 8, #happens in at least 5% of the 37 observations
                                  link = "unweighted",
                                  min.prob = 0.05)

complot(FF_min_all_net, title = "Female-female")

ggsave("FF_complot.png", device = "png", path = "Paper_1/figures/", width = 10, height = 6)

rm(FF_min_all_greet, FF_min_all_net, MM_min_all_greet, MM_min_all_net, MM_subset, FF_subset)
# remove excess objects from environment 

### Female-male ################

set.seed(55)
#to look at only female-male
FM_subset <- min_DF %>%
  cbind(video_info) %>%
  filter(Greeting_type == "female male") %>%
  select(c(1:25))


FM_min_all_greet<- netfacs(
  data = FM_subset,
  condition = NULL,
  ran.trials = 2000,
  combination.size = 3,
  use_parallel = TRUE
)

FM_min_all_net <- netfacs_network(netfacs.data = FM_min_all_greet,
                                  significance = 0.05, # cut off value of 0.05 due to low sample size
                                  min.count = 4, #happens in at least 5% of the 37 observations
                                  link = "unweighted",
                                  min.prob = 0.05)

complot(FM_min_all_net, title = "Female-male")


ggsave("FM_complot.png", device = "png", path = "Paper_1/figures/", width = 10, height = 6)

rm(FM_min_all_net, FM_min_all_greet, FM_subset)


### Male-female ################

set.seed(55)
#to look at only male-female
MF_subset <- min_DF %>%
  cbind(video_info) %>%
  filter(Greeting_type == "male female") %>%
  select(c(1:25))


MF_min_all_greet<- netfacs(
  data = MF_subset,
  condition = NULL,
  ran.trials = 2000,
  combination.size = 3,
  use_parallel = TRUE
)

MF_min_all_net <- netfacs_network(netfacs.data = MF_min_all_greet,
                                  significance = 0.05, # cut off value of 0.05 due to low sample size
                                  min.count = 3, #happens in at least 5% of the 37 observations
                                  link = "unweighted",
                                  min.prob = 0.05)

complot(MF_min_all_net, title = "Male-female")


ggsave("MF_complot.png", device = "png", path = "Paper_1/figures/", width = 10, height = 6)

rm(MF_min_all_greet, MF_min_all_net, MF_subset) #remove excess objects from environment 


# 3.3 APPROACHER SEX ############################

## 3.3.1 Male vs female ########################
set.seed(22) # set seed for each analysis so they can run sep and be reproducible

male_greeters<- netfacs(
  data = max_DF, # this is the data matrix
  condition = video_info$Greeter_sex, # info about condition of each case
  test.condition = "male", # condition we are interested in
  null.condition = "female", # null condition
  duration = NULL, 
  ran.trials = 1000, # number of randomizations
  control = list(video_info$Greetee_sex, video_info$other_infant), # control variables
  random.level = c(video_info$Greeter), # Works like a random effect.
  combination.size = 3, # limit the analysis
  tail = "upper.tail", # p-value reflects upper-tail
  use_parallel = T, # use parallel processing
  n_cores = NULL # number of cores for parallel processing
)


## Select the types that we are interested in

test_behvs_M <- filter(male_greeters$result, combination.size == 1, combination %in% c("EarFlattening", "LipSmack", "Mount", "Present", "HqContact")) %>%
  select(-c(2))

test_behvs_M

# Upsample to get specificity
specificity(male_greeters, upsample = TRUE)


saveRDS(test_behvs_M, "Paper_1/figures/male-outcome-table-5.rds")


### Element figure ################

MLS_plot <- distribution.plot(netfacs.data = male_greeters)$"LipSmack" +  #creating the figure for lipsmacking in males
  xlab("Probability") +
  ggtitle("Lip-Smacking") +
  theme(
    axis.text.x = element_text(size = 10),
    axis.text.y = element_text(size = 10),  
    axis.title.x = element_text(size = 10),
    axis.title.y = element_text(size = 10),
    plot.title = element_text(size = 11),
    panel.grid.major = element_blank(), 
    panel.grid.minor = element_blank(),
    panel.background = element_blank(), 
    axis.line = element_line(colour = "black"))+
  scale_x_continuous(limits = c(0,0.6))

MP_plot <- distribution.plot(netfacs.data = male_greeters)$"Present" + # presenting in males
  xlab("Probability") +
  theme(
    axis.text.x = element_text(size = 10),
    axis.text.y = element_text(size = 10),  
    axis.title.x = element_text(size = 10),
    axis.title.y = element_text(size = 10),
    plot.title = element_text(size = 11),
    panel.grid.major = element_blank(), 
    panel.grid.minor = element_blank(),
    panel.background = element_blank(), 
    axis.line = element_line(colour = "black"))+
  scale_x_continuous(limits = c(0,0.6)) 

MEF_plot <- distribution.plot(netfacs.data = male_greeters)$"EarFlattening" + #ear flattening in males
  xlab("Probability")+
  ggtitle("Ear-Flattening")+
  theme(
    axis.text.x = element_text(size = 10),
    axis.text.y = element_text(size = 10),  
    axis.title.x = element_text(size = 10),
    axis.title.y = element_text(size = 10),
    plot.title = element_text(size = 11),    
    panel.grid.major = element_blank(), 
    panel.grid.minor = element_blank(),
    panel.background = element_blank(), 
    axis.line = element_line(colour = "black"))+
  scale_x_continuous(limits = c(0,0.6)) 

MHQ_plot <- distribution.plot(netfacs.data = male_greeters)$"HqContact" + # hindquarter contact in males
  xlab("Probability")+
  ggtitle("Hind-quarter Contact")+
  theme(
    axis.text.x = element_text(size = 10),
    axis.text.y = element_text(size = 10),  
    axis.title.x = element_text(size = 10),
    axis.title.y = element_text(size = 10),
    plot.title = element_text(size = 11),    
    panel.grid.major = element_blank(), 
    panel.grid.minor = element_blank(),
    panel.background = element_blank(), 
    axis.line = element_line(colour = "black"))+
  scale_x_continuous(limits = c(0,0.6))

# These plots are all used below with the female versions to create one main plot

# extract that looks at any behaviour combo that is significant with a probability increase over one that includes one of our behaviours of interest
male_fac_extracts <- filter(male_greeters$result, (pvalue <= 0.05) & (prob.increase > 1) & (count >= 4) & str_detect(combination, "EarFlattening|Mount|HqContact|Present|LipSmack")) 

male_fac_extracts

saveRDS(male_fac_extracts, "Paper_1/figures/male-all-table.rds")


## 3.3.1 Female vs male ########################


female_greeters<- netfacs(
  data = max_DF, # this is the data matrix
  condition = video_info$Greeter_sex, # info about condition of each case
  test.condition = "female", # condition we are interested in
  null.condition = "male", # null condition
  duration = NULL,
  ran.trials = 1000, # number of randomizations
  control = list(video_info$Greetee_sex, video_info$other_infant), # control variables
  #random.level = c(video_info$Greeter), # Attempted to include random effect of greeter, but would not converge/run successfully so removed.
  combination.size = 3, # limit the analysis to make things faster,
  tail = "upper.tail", # p-value from upper-tail
  use_parallel = T, 
  n_cores = NULL 
)


#extract results on our 5 behaviours of interest
test_behvs_F <- filter(female_greeters$result, combination.size == 1, combination %in% c("EarFlattening", "LipSmack", "Mount", "Present", "HqContact")) %>%
  select(-c(2))

test_behvs_F 

# upsample for specificity 
spec_fem_greeters <- specificity(female_greeters, upsample = TRUE)

saveRDS(test_behvs_F, "Paper_1/figures/female-outcome-table-5.rds")


#extract any results relating to combinations which include at least one of our behaviours of interest
female_fac_extracts <- filter(female_greeters$result, (pvalue <= 0.05) & (prob.increase > 1) & (count >= 4) & str_detect(combination, "EarFlattening|Mount|HqContact|Present|LipSmack")) 

female_fac_extracts

saveRDS(female_fac_extracts, "Paper_1/figures/female-all-table.rds")


### Element figure ##############
#creating plots for each of the 4 behaviours of interest

FLS_plot <- distribution.plot(netfacs.data = female_greeters)$"LipSmack" + # female lipsmacking plot
  xlab("Probability") +
  ggtitle("Lip-Smacking") +
  theme(
    axis.text.x = element_text(size = 10),
    axis.text.y = element_text(size = 10),  
    axis.title.x = element_text(size = 10),
    axis.title.y = element_text(size = 10),
    plot.title = element_text(size = 11),
    panel.grid.major = element_blank(), 
    panel.grid.minor = element_blank(),
    panel.background = element_blank(), 
    axis.line = element_line(colour = "black"))+
  scale_x_continuous(limits = c(0,0.6))

FP_plot <- distribution.plot(netfacs.data = female_greeters)$"Present" + # female presenting plot
  xlab("Probability") +
  theme(
    axis.text.x = element_text(size = 10),
    axis.text.y = element_text(size = 10),  
    axis.title.x = element_text(size = 10),
    axis.title.y = element_text(size = 10),
    plot.title = element_text(size = 11),
    panel.grid.major = element_blank(), 
    panel.grid.minor = element_blank(),
    panel.background = element_blank(), 
    axis.line = element_line(colour = "black"))+
  scale_x_continuous(limits = c(0,0.6)) 

FEF_plot <- distribution.plot(netfacs.data = female_greeters)$"EarFlattening" + #female ear flattening plot
  xlab("Probability")+
  ggtitle("Ear-Flattening")+
  theme(
    axis.text.x = element_text(size = 10),
    axis.text.y = element_text(size = 10),  
    axis.title.x = element_text(size = 10),
    axis.title.y = element_text(size = 10),
    plot.title = element_text(size = 11),    
    panel.grid.major = element_blank(), 
    panel.grid.minor = element_blank(),
    panel.background = element_blank(), 
    axis.line = element_line(colour = "black"))+
  scale_x_continuous(limits = c(0,0.6)) 

FHQ_plot <- distribution.plot(netfacs.data = female_greeters)$"HqContact" + # female hindquarter contact plot
  xlab("Probability")+
  ggtitle("Hind-quarter Contact")+
  theme(
    axis.text.x = element_text(size = 10),
    axis.text.y = element_text(size = 10),  
    axis.title.x = element_text(size = 10),
    axis.title.y = element_text(size = 10),
    plot.title = element_text(size = 11),    
    panel.grid.major = element_blank(), 
    panel.grid.minor = element_blank(),
    panel.background = element_blank(), 
    axis.line = element_line(colour = "black"))+
  scale_x_continuous(limits = c(0,0.6))

## Figure 3.2 ########################

# create the whole figure, putting together the male and female plots
figure <- (MLS_plot / MP_plot / MEF_plot / MHQ_plot) & scale_y_continuous(
  limits = c(0, 42))  | (FLS_plot / FP_plot / FEF_plot / FHQ_plot) & scale_y_continuous(limits = c(0, 42)) 

figure + plot_annotation(
    title = "Observed vs expected probability in male (left) and female (right) approachers") # add a title to the plot

ggsave("probability_figure.png", plot = print(figure + plot_annotation(
  title = "Observed vs expected probability in male (left) and female (right) approachers"
)), device = "png", path = "Paper_1/figures/", width = 8, height = 6, dpi = 600)   # save the plot

# remove unnecessary objects from environment 
rm(FEF_plot, female_fac_extracts, FHQ_plot, figure, FLS_plot, FP_plot, male_fac_extracts, 
   MEF_plot, MHQ_plot, MLS_plot, MP_plot, test_behvs_F, test_behvs_M)


## 3.3.2 Across sex combinations ###########################
###MM vs all ##################
# looking at behaviours that happen in male-male greetings more than the other types

MM <- netfacs(
  data = max_DF, # this is the data matrix
  condition = video_info$Greeting_type, # info about condition of each case
  test.condition = "male male", # condition we are interested in
  null.condition = NULL, # null condition
  duration = NULL,
  ran.trials = 1000, # number of randomizations
  control = list(video_info$other_infant), # control variables
  random.level = c(video_info$Greeter), # random effect.
  combination.size = 3, # limit the analysis to make things faster,
  tail = "upper.tail", # upper-tail p-value
  use_parallel = T, 
  n_cores = NULL 
)


# extract that looks at any behaviour combo that is significant with a probability increase over one that includes one of our behaviours of interest

MM_extracts <- filter(MM$result, (pvalue <= 0.05) & (prob.increase > 1) & (count >= 4) & str_detect(combination, "EarFlattening|Mount|HqContact|Present|LipSmack")) 

MM_extracts

#upsample for specificity and then filter to include only behaviours we are interested in
MM_spec <- specificity(MM, upsample = TRUE)
MM_spec %>% 
  filter((pvalue <= 0.05) & (prob.increase > 1) & (count >= 4) & str_detect(combination, "EarFlattening|Mount|HqContact|Present|LipSmack")) %>%
  select(1,2, specificity)


### FF vs all ###########################
# run bootstrapping
FF <- netfacs(
  data = max_DF, # this is the data matrix
  condition = video_info$Greeting_type, # info about condition of each case
  test.condition = "female female", # condition we are interested in
  null.condition = NULL, # null condition
  duration = NULL, 
  ran.trials = 1000, # number of randomizations
  control = list(video_info$other_infant), # control variables
  random.level = c(video_info$Greeter), # random effect.
  combination.size = 3, # limit the analysis to make things faster,
  tail = "upper.tail", # upper-tail p-value
  use_parallel = T,
  n_cores = NULL
)

# extract that looks at any behaviour combo that is significant with a probability increase over one that includes one of our behaviours of interest
FF_extracts <- filter(FF$result, (pvalue <= 0.05) & (prob.increase > 1) & (count >= 4) & str_detect(combination, "EarFlattening|Mount|HqContact|Present|LipSmack")) 

FF_extracts

#upsample to get specificity
FF_spec <- specificity(FF, upsample = TRUE)

FF_spec %>% #filter so we only get what we are interested in
  filter((pvalue <= 0.05) & (prob.increase > 1) & (count >= 4) & str_detect(combination, "EarFlattening|Mount|HqContact|Present|LipSmack")) %>%
  select(1,2, specificity)

saveRDS(FF_extracts, "Paper_1/figures/FF-all-table.rds")
write_csv(FF_extracts, "Paper_1/processed-data/FF_all_table.csv")


### MF vs all ######################
# run bootstrapping
MF <- netfacs(
  data = max_DF, # this is the data matrix
  condition = video_info$Greeting_type, # info about condition of each case
  test.condition = "male female", # condition we are interested in
  null.condition = NULL, # null condition
  duration = NULL, 
  ran.trials = 1000, # number of randomizations
  control = list(video_info$other_infant), # control variables
  random.level = c(video_info$Greeter), # random effect.
  combination.size = 3, # limit the analysis to make things faster,
  tail = "upper.tail", # upper-tail p-value
  use_parallel = T, 
  n_cores = NULL 
)


# extract that looks at any behaviour combo that is significant with a probability increase over one that includes one of our behaviours of interest
MF_extracts <- filter(MF$result, (pvalue <= 0.05) & (prob.increase > 1) & (count >= 4) & str_detect(combination, "EarFlattening|Mount|HqContact|Present|LipSmack")) 

MF_extracts

#upsample for specificity 
MF_spec <- specificity(MF, upsample = TRUE)

MF_spec %>% #filter so we're only looking at what we're interested in
  filter((pvalue <= 0.05) & (prob.increase > 1) & (count >= 4) & str_detect(combination, "EarFlattening|Mount|HqContact|Present|LipSmack")) %>%
  select(1,2, specificity)

### FM vs all ######################### 
# run bootstrapping
FM <- netfacs(
  data = max_DF, # this is the data matrix
  condition = video_info$Greeting_type, # info about condition of each case
  test.condition = "female male", # condition we are interested in
  null.condition = NULL, # null condition
  duration = NULL, 
  ran.trials = 1000, # number of randomizations
  control = list(video_info$other_infant), # control variables
  random.level = c(video_info$Greeter), # random effect.
  combination.size = 3, # limit the analysis to make things faster,
  tail = "upper.tail", # upper-tail p-value
  use_parallel = T, 
  n_cores = NULL 
)

# extract that looks at any behaviour combo that is significant with a probability increase over one that includes one of our behaviours of interest
FM_extracts <- filter(FM$result, (pvalue <= 0.05) & (prob.increase > 1) & (count >= 4) & str_detect(combination, "EarFlattening|Mount|HqContact|Present|LipSmack")) 

FM_extracts

#upsample for specificity 
FM_spec <- specificity(FM, upsample = TRUE)

FM_spec %>% # filter so we are only looking at what we are interested in
  filter((pvalue <= 0.05) & (prob.increase > 1) & (count >= 4) & str_detect(combination, "EarFlattening|Mount|HqContact|Present|LipSmack")) %>%
  select(1,2, specificity)

# simplified comparison lists

# MM size 1
netfacs.extract(MM, level = 1, min.count = 4, min.prob = 0, significance = 0.01)

# FF size 1
netfacs.extract(FF, level = 1, min.count = 4, min.prob = 0, significance = 0.01)

# MM size 2
netfacs.extract(MM, level = 2, min.count = 4, min.prob = 0, significance = 0.01)

## FF size 2
netfacs.extract(FF, level = 2, min.count = 4, min.prob = 0, significance = 0.01)

# FM size 1
netfacs.extract(FM, level = 1, min.count = 4, min.prob = 0, significance = 0.01)

# MF size 1
netfacs.extract(MF, level = 1, min.count = 4, min.prob = 0, significance = 0.01)

#FM size 2
netfacs.extract(FM, level = 2, min.count = 4, min.prob = 0, significance = 0.01)

# MF size 2
netfacs.extract(MF, level = 2, min.count = 4, min.prob = 0, significance = 0.01)


# CLEAN UP #################################################

# Clear environment
rm(list = ls()) 

# Clear packages
pacman::p_unload(negate=TRUE)  # Remove all add-ons

# Clear plots
dev.off()  # But only if there IS a plot

# Clear console
cat("\014")  # ctrl+L

# Clear mind :)
```
